# Salinity induces discontinuous protoxylem by a DELLA-dependent mechanism promoting salt tolerance in Arabidopsis seedlings

**DOI:** 10.1101/2022.02.13.480244

**Authors:** Frauke Augstein, Annelie Carlsbecker

## Abstract

- Salinity is detrimental to plants and developmental adjustments limiting salt uptake and transport is therefore important for acclimation to high salt. These parameters may be influenced by xylem morphology, however, how plant root xylem development is affected by salt stress remains unclear.
- Through detailed phenotypic analyses, molecular and genetic techniques, we demonstrate that salt causes distinct effects on Arabidopsis seedling root xylem, and reveal underlying molecular mechanisms.
- Salinity causes intermittent inhibition of protoxylem cell differentiation, generating protoxylem gaps, in Arabidopsis and other eudicot seedlings. The extent of protoxylem gaps positively correlates with salt tolerance. Reduced gibberellin signaling is required for protoxylem gap formation. Mutant analyses reveal that the xylem differentiation regulator VASCULAR RELATED NAC DOMAIN 6 (VND6), along with secondary cell wall-producing and cell wall modifying enzymes, including EXPANSIN A1 (EXP1), are involved in protoxylem gap formation, in a DELLA-dependent manner.
- Salt stress impacts seedling survival and formation of protoxylem gaps is a means of enhancing salt tolerance. Salt stress likely reduces levels of bioactive gibberellins, stabilizing DELLAs, which in turn activate multiple factors modifying protoxylem differentiation. Formation of protoxylem gaps is induced in diverse eudicot species suggesting that this is an evolutionary conserved response for salt acclimation in seedlings.

## Introduction

Survival of plant seedlings is affected by many environmental parameters such as available water and soil salinity. Salt has a negative impact on the plant both through its osmotic effect, which may result in reduced ability for water uptake, and because of ion toxicity (Munns and Tester, 2008). It affects many important processes including photosynthesis, respiration, ion uptake and membrane integrity (West et al., 2004; Tavakkoli et al., 2011; Talei et al., 2012; Mansour, 2013; Awlia et al., 2021; Zhao et al., 2021). Salt stress tolerance is expected to involve both avoidance mechanisms and reduced uptake and transport of salt ions (Møller et al., 2009). The initial response to saline conditions is a growth arrest of both primary and lateral roots followed by a temporally dynamic acclimation process in which growth is restored and salt tolerance mechanisms activated (Geng et al., 2013). Thus, it is conceivable that salt stress also affects development of the water transporting tissue, the xylem, as that would impact salt uptake. However, how salt affects xylem development is not well known.

The xylem harbors vessel strands of hollow cells reinforced with lignified secondary cell walls (SCW). In the Arabidopsis root the xylem forms an axis traversing the stele. The two outer strands of the xylem axis differentiate as protoxylem with annular or helical SCW, while metaxylem with pitted SCW occupy the central positions of the axis (Fig. 1a). The diameter and shape of the SCWs are thought to influence hydraulic conductance, and thus the xylem shape correlates with drought resistance in many different species (Arend and Fromm, 2007; Awad et al., 2010; Tang et al., 2018; Yu et al., 2021). Recently, we and others showed that xylem formation is plastic and responds to water availability. Under conditions of reduced water availability, extra protoxylem strands form, and metaxylem differentiates closer to the root tip (Ramachandran et al., 2018; Bloch et al., 2019; Ramachandran et al., 2021). The hormone abscisic acid (ABA) mediates these developmental responses by at least two different mechanisms. Firstly, ABA promotes the production of miR165 in the endodermis (Ramachandran et al., 2018; Bloch et al., 2019; Ramachandran et al., 2021). This miRNA moves into the stele to reduce homeodomain leucine zipper class III (HD-ZIP III) mRNA levels eventually leading to protoxylem formation in place of metaxylem (Carlsbecker et al., 2010; Miyashima et al., 2011). Secondly, ABA directly promotes expression of VASCULAR RELATED NAC-DOMAIN (VND) transcription factors within the immature xylem cells, where VND7 promotes protoxylem formation, and VND2 and VND3 metaxylem differentiation closer to the root tip (Ramachandran et al., 2021).

**Fig. 1:**
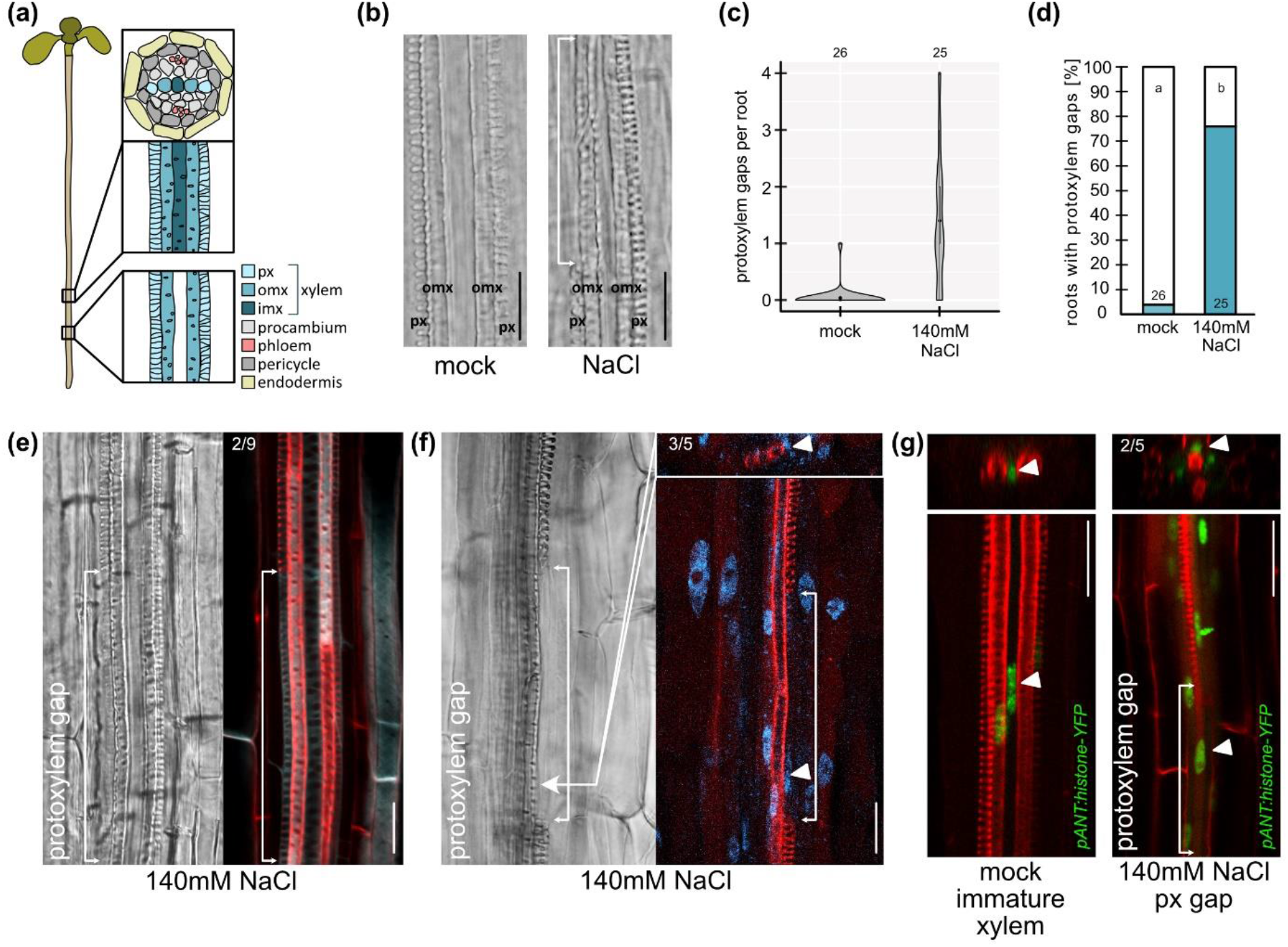
Protoxylem gaps are formed in response to salt. All images depict roots of 6-day old Arabidopsis seedlings grown for 3 days on 140mM NaCl or under mock conditions. **(a)** Cartoon of Arabidopsis seedling root xylem in longitudinal view and stele + endodermis in cross section. px, protoxylem; omx, outer metaxylem; imx, inner metaxylem. **(b)** Differential interference (DIC) images of root xylem. White arrow indicates a protoxylem gap. Scale bars, 50 μm. **(c)** Quantification of protoxylem gaps per root. Number of analyzed roots (n) are indicated above the graph. **(d)** Quantification of roots exhibiting protoxylem gaps. n is indicated on the bars; letters indicate statistical significance with multiple Fisher’s exact test and BH correction, p < 0.05. **(e - g)** DIC and confocal micrographs of root xylem. White arrows indicate protoxylem gaps. Arrowheads point at nuclei in protoxylem gap cells. Numbers indicate fraction of analyzed gaps that showed **(e)** cellulose SCW pattern, **(f)** nuclear signal within the gap, **(g)** *pANT:histone-YFP* expression within the gap. Turquoise, Calcofluor White staining cellulose; Red, Basic Fuchsin staining lignin; Blue, DAPI staining nucleus; Green, *pANT:histone-YFP*. Scale bars, 20 μm.

Salt stress also triggers ABA signaling, but recent research has in addition highlighted the importance of gibberellin (GA) levels and signaling. Reduced GA levels or signaling may result in salt stress tolerance (Colebrook et al., 2014), while the absence of the DELLA repressors of GA-signaling make Arabidopsis less salt tolerant (Achard et al., 2006). In line with these findings, salt stress leads to a reduction in bioactive GA levels, which in turn stabilizes DELLAs (Achard et al., 2006; Magome et al., 2008). Under normal conditions, GAs affect xylem lignification in both primary and secondary development in several different species (Eriksson et al., 2000; Mauriat and Moritz, 2009; Gou et al., 2011; Ragni et al., 2011; Guo et al., 2015; Wang et al., 2017; Singh et al., 2019), GAs promote xylem formation in secondary development and are important for fiber development (Mauriat and Moritz, 2009; Ragni et al., 2011; Felipo-Benavent et al., 2018). Furthermore, DELLAs are implicated in the regulation of cell wall synthesis and remodeling in Arabidopsis (Locascio et al., 2013).

Here, we analyze the effect of salt stress on Arabidopsis seedling root xylem development. We show that salt stress results in discontinuous differentiation of the protoxylem strands generating protoxylem gaps. Mutants with enhanced protoxylem gap formation display enhanced tolerance to salt stress, while those with inhibited gap formation were more salt sensitive, suggesting that protoxylem gaps promote salt stress tolerance. We also show that formation of protoxylem gaps under salt stress requires DELLA-mediated repression of GA signaling. Under salt, DELLAs promote VND6 and associated SCW enzymes as well as multiple cell wall modifying enzymes including alpha-expansins such as EXP1 also called EXPA1. The loss of *VND6* or *EXP1* consequently results in less protoxylem gaps forming under salt stress.

## Material and Methods

### Plant material and growth conditions

Seeds were surface sterilized using 70% ethanol for 20 min and 95% ethanol for 2 min, and then rinsed four times for 2 min in sterile water. The seeds were plated on 0.5x Murashige and Skoog (MS) medium (Murashige and Skoog, 1962), pH 5.7-5.8, with 0.05% MES monohydrate and 1% Bactoagar, and stratified for 48 h at 4°C. For all experiments, plants were grown vertically on 25 mm pore Sefar Nitex 03-25/19 mesh, and transferred to new plates by transferring the mesh with the plants on for minimal disturbance. For GA3, a stock solution in 99.9% EtOH was prepared; for GA4+7 and PAC stock solutions were prepared in DMSO. For plates with NaCl, a 3M stock solution was used and diluted in the medium to the indicated concentration. Mannitol was added directly into the medium after autoclaving. For Arabidopsis xylem phenotyping experiments, 3-day old seedlings were transferred to treatment-plates for 3 days. For tolerance assays, 3-day-old plants were left on NaCl-plates for 4 to 7 days. For RNA-sequencing analysis, 5-day old seedlings were used and exposed to salt for the indicated time. All material was collected at the same time, in the afternoon, to avoid circadian clock effects. For phenotyping of other species than Arabidopsis, seedlings were grown until roots reached approximately 1cm in length before transfer to salt for 3 days. Detailed information about all genotypes can be found in Table S2.

### Root length measurements

Root lengths were measured using Fiji/Image J.

### Xylem phenotype analysis

For analysis of xylem morphology, roots were mounted in chloralhydrate solution, 8:2:1 chloralhydrate:glycerol:water (w/v/v), and visualized as previously described using a Zeiss Axioscope A1 microscope at 40X magnification with differential interference contrast (DIC) optics (Ramachandran et al., 2018; Ramachandran et al., 2021). Frequency of plants exhibiting the protoxylem gap phenotype was scored, and number of gaps per root was counted.

### Salt tolerance assay

Coloring of cotyledons of seedlings grown on high salt was determined after 4 or 7 days. Plants exhibiting white, pale green or green cotyledons were counted separately. From the fraction of each category a survival score was calculated multiplying white with 1, pale with 3 and green with 5 divided by the sum, following (Gibbs et al., 2011). Here data is shown from one experiment with three to five replicates per genotype-treatment combination. The experiment was repeated two to three times (Table S3)

### Confocal analysis

For parallel staining with Basic Fuchsin and Calcofluor White or DAPI, we followed a modified fixation protocol from (Ursache et al., 2018). 6-day old seedlings were fixed with 4% PFA in 1xPBS, for 1h for Basic Fuchsin and Calcofluor White, 10-15 min for Basic Fuchsin and DAPI staining or 10-15 min for Basic Fuchsin stain of transcriptional reporter lines. This was followed by two times washing for 1 min with 1xPBS, and then clearing over night with ClearSee (10% xylitol, 15% sodium deoxycholate, 25% urea in water). For the Basic Fuchsin stain, seedlings were then stained with 0.2% Basic Fuchsin (in ClearSee) over night, and washed with ClearSee two times. For Calcofluor White staining, seedlings were stained with 0.1% Calcofluor White for 30 min, and washed with ClearSee for 30 min. For visualization with confocal microscope roots were mounted directly in ClearSee. For DAPI stain, seedlings were mounted in 0.4μl of 5mg/ml DAPI solution in 10 ml H_2_O and visualized directly. For *RGA:GFP-RGA* reporter analysis, roots were mounted in 40mM propidium iodide (PI) solution between two coverslips and imaged immediately.

Confocal micrographs were captured using Zeiss LSM780 inverted Axio Observer with supersensitive GaAsP detectors or an LSM800 inverted Confocal microscope. For Calcofluor White a 405 nm laser was used for excitation and emission wavelengths of 410-475 nm were captured. For Basic Fuchsin images, 561 nm excitation and 600-700 nm emission. For DAPI, 405 nm excitation and 410-511 emission. For reporter lines expressing GFP and stained with PI, 561nm excitation and 650-700 nm emission were used for PI and 488 nm excitation and 410-523 nm emission for GFP. For reporter lines expressing YFP, 514 nm excitation and 518-544 emission was used. For quantification of fluorescence intensity all imaging parameters were kept the same when imaging mock and treated roots. Fluorescence intensity was quantified using CellSeT (Pound et al., 2012).

### RNA sequencing analysis

5-day old Arabidopsis seedlings of Landsberg *erecta* (wildtype), *gai-t6 rga-t2 rgl1-1 rgl2-1 rgl3-4* (della5x), were grown on 140mM NaCl or under mock conditions for 1h or 8h. For the 8h timepoint *ga4* and *gai* were grown in parallel. Three biological replicates, each consisting of 50-100 seedlings, were collected for each treatment-genotype combination. The lower part of the root (1 cm) was collected directly in RLT buffer (QIAGEN) and frozen in liquid nitrogen. RNA was extracted using the RNeasy Plant Mini Kit (QIAGEN). RNA concentration was measured with Qubit BR RNA Assay and quality and integrity of the RNA was analyzed with the Agilent Bioanalyzer 2100 system (Agilent Technologies, CA, USA). A total amount of 1000 ng RNA per sample was used for library preparation. Library preparation and sequencing was performed by Novogene (UK) on their Illumina sequencing platform with paired-end read length of 150 and 250-300bp cDNA library resulting in 5.9 to 8.3 G raw data per sample (241.9 G total). Fastp was used for quality assessment and adapter trimming (Chen et al., 2018). Mapping to the *Arabidopsis thaliana* reference genome (TAIR10) was done using Hisat2 (Kim et al., 2019). 96% of the total reads were mapped to the Arabidopsis genome, whereby 93% of the total reads were uniquely mapped. Count files were generated using HTSeq-Count (Anders et al., 2015). Differential expression analysis was done using DESeq2 in Bioconductor (Huber et al., 2015). For statistical analysis of NaCl effects on the different genotypes compared to wildtype, a DESeq2 model including a combinatorial effect was used (~genotype+genotype:condition). Log2 fold changes (FC) were extracted from the pairwise comparison of mock versus treatment for each genotype, while p values and adjusted p values were extracted from the comparison between the mutants and wildtype. The effect of the different genotypes under mock condition was analyzed in a separate differential expression analysis and all values were extracted from the pairwise comparison of wildtype versus mutant. A cut-off > 0.5 or < −0.5 log2 FC was applied.

GO-term analysis was performed with PANTHER Classification System (Mi et al., 2019; Mi et al., 2021) using *Arabidopsis thaliana* as a reference and the GO annotation data set “biological process complete”. Fisher’s Exact test was selected as test type and Bonferroni correction for multiple testing was performed. GO-term clustering was performed using Revigo using p-values (Supek et al., 2011). *Arabidopsis thaliana* was used as a reference, obsolete GO terms were removed and SimRel was used as semantic similarity measure.

### Statistical analysis

For categorical data, Fisher’s exact test using the fisher.multcomp() function of the ‘RVAideMemoire’ package (Hervé, 2021) in R was performed and p values less than 0.05 were considered significant. The p-values were corrected for multiple testing using the Benjamini and Hochberg correction (Benjamini and Hochberg, 1995). For other data, Two-way ANOVA, using the aov() function in R combined with a Tukey post hoc test or t-tests using the t.test() function were used as indicated. Statistical tests and significance thresholds are mentioned in figure legends. Number of roots analyzed are mentioned in the corresponding figures.

## Results

### Salt stress inhibits local protoxylem differentiation causing discontinuous xylem strands

To assess how salt stress affects seedling root xylem development we grew 3-day-old Arabidopsis Col-0 seedlings on 140 mM NaCl, which is a high but non-lethal concentration (Dinneny et al., 2008), for 3 days, and then analyzed primary root growth and xylem morphology. Consistent with previous findings, where transfer to salt resulted in an initial growth inhibition relieved after some time of acclimation (Geng et al., 2013), root growth on salt was substantially reduced (Fig. S1a). We found that the predominant effect on xylem morphology was an appearance of discontinuous protoxylem, seen as protoxylem gaps spread along the part of the xylem, which had differentiated during growth on high salt (Fig. 1b, S1b). Hence, the protoxylem gaps were likely not an effect of an initial root growth inhibition upon transfer to high salt concentrations, but rather an effect of the salt stress itself. In 75% of the seedlings analyzed, up to four protoxylem gaps on either one or both xylem strands were induced, with an average of 1.4 gaps per root (Fig. 1c, d). Growth for 3 days on concentrations ranging from 80 to 140 mM of NaCl revealed a concentration dependence in the frequency of plants displaying protoxylem gaps (Fig. S1C). Growth on 280 mM mannitol, iso-osmolaric to 140 mM NaCl, did not result in protoxylem gap formation (Fig. S1C) suggesting that xylem gap formation is linked to the ionic stress rather than the osmotic stress.

While the formation of xylem gaps was consistently observed in response to salt stress, it is possible that this is merely a symptom of the toxicity of the salt ions in combination with the osmotic stress causing collapsed cells rather than a developmentally controlled response suppressing local xylem differentiation. To distinguish between these possibilities, we analyzed xylem strands double stained with Basic Fuchsin and Calcofluor White to visualize lignin and cellulose (Ursache et al., 2018). This revealed that while lignin was absent in the gaps, cellulose was detected. In 2 of 9 gaps we observed cellulose with SCW patterns (Fig. 1e) reminiscent of the fully lignified neighboring cells, while other gaps displayed a thin cell wall, indicating that these cells only had a primary cell wall. This suggests that salt impacted on SCW formation and lignin deposition. Following SCW formation a xylem cell would undergo programmed cell death (PCD) to form a hollow tube for water transport (Schuetz et al., 2013). To analyze if the gap cells maintained a nucleus we fixed and stained salt-grown roots with DAPI for DNA and Basic Fuchsin for lignin. In the lignified xylem cells surrounding a gap we could not detect DAPI-staining, indicating that these cells had undergone PCD. However, in 3 of 5 non-lignified xylem gap cells, we detected localised DAPI-staining indicating an intact nucleus (Fig. 1f). Furthermore, *pANT:histone-YFP*, a reporter for *AINTEGUMENTA* active in procambium, immature xylem cells and vascular cambium (Randall et al., 2015), was expressed in the salt-induced gap cells, suggesting that these cells displayed procambial or immature xylem cell identity (Fig. 1g). As certain xylem gap cells apparently were living, we tested if gap cells could resume differentiation if the plants grown on salt were allowed to continue growth on normal medium. After transfer to normal conditions, we observed similar number of gaps in the root section previously grown under high salt conditions, whereas no additional xylem gaps were formed (Fig. S1d). Hence, our analyses suggest that salt triggers a local non-reversible suppression of xylem cell differentiation.

### Xylem gap formation is not ABA mediated

Extended growth on 140 mM NaCl for 3, 5 or 7 days revealed that the formation of xylem gaps is a relatively early response, followed by formation of additional protoxylem strands (Fig. S1e, f, g). Additional protoxylem formation also occurs under water deficiency or after treatment with ABA (Ramachandran et al., 2018; Bloch et al., 2019). To test if ABA signalling is similarly important for the generation of xylem gaps we made use of the dominant negative *abi1-1* mutant, in which ABA signalling is suppressed (Leung et al., 1994; Meyer et al., 1994). As previously found, *abi1-1* reduced the frequency of additional protoxylem formed upon osmotic stress (Fig. S1h), and it also partially suppressed the formation of metaxylem closer to the root tip which happens under growth on both salt and on mannitol (Fig. S1i, j; Ramachandran et al., 2018; Ramachandran et al., 2021). In contrast, protoxylem gap formation was not affected by *abi1-1* (Fig. S1k), nor did ABA-treatment cause xylem gaps (Fig. S1l). Hence, while formation of additional protoxylem and earlier metaxylem differentiation are ABA-mediated effects also under growth on salt, xylem gap formation does not require ABA signalling.

### Salt stress induces root xylem gaps in several eudicot species

To elucidate if high salinity-induced protoxylem gaps are a general phenomenon in eudicot seedlings, we studied three additional species. We selected tomato (*Solanum lycopersicum*; Solanaceae, subclass Asteridae) which like Arabidopsis (Brassicacacea, subclass Rosidae) is considered moderately salt sensitive (Munns and Tester, 2008; Sun et al., 2010), sugar beet (*Beta vulgaris*; Amaranthaceae, subclass Rosidae) which is salt tolerant (Skorupa et al., 2019), and the halophyte *Eutrema salsugineum* (Brassicaceae) (Yang et al., 2013). Similar to Arabidopsis, tomato seedlings formed gaps in at least one of the protoxylem strands in most roots analyzed when grown on 140 mM NaCl (Fig. 2a, d). When grown on normal media a relatively high percentage of sugar beet seedling roots displayed xylem gaps, and while 140 mM NaCl did not significantly enhance gap formation, growth on 200 mM NaCl did (Fig. 2b, e, S2a). In contrast to sugar beet, no protoxylem gaps were observed in *E. salsugineum* on normal media, but like sugar beet 200 mM NaCl triggered a substantial increase in xylem gaps, while 140mM did not (Fig. 2c, f, S2b). Growth on 200mM salt also induced additional protoxylem strand formation (Fig. S2c). These results indicate that xylem gaps form upon salt stress in distantly related eudicot species, regardless if these are salt sensitive, salt tolerant or even halophytes, if exposed to high enough salt concentration.

**Fig. 2:**
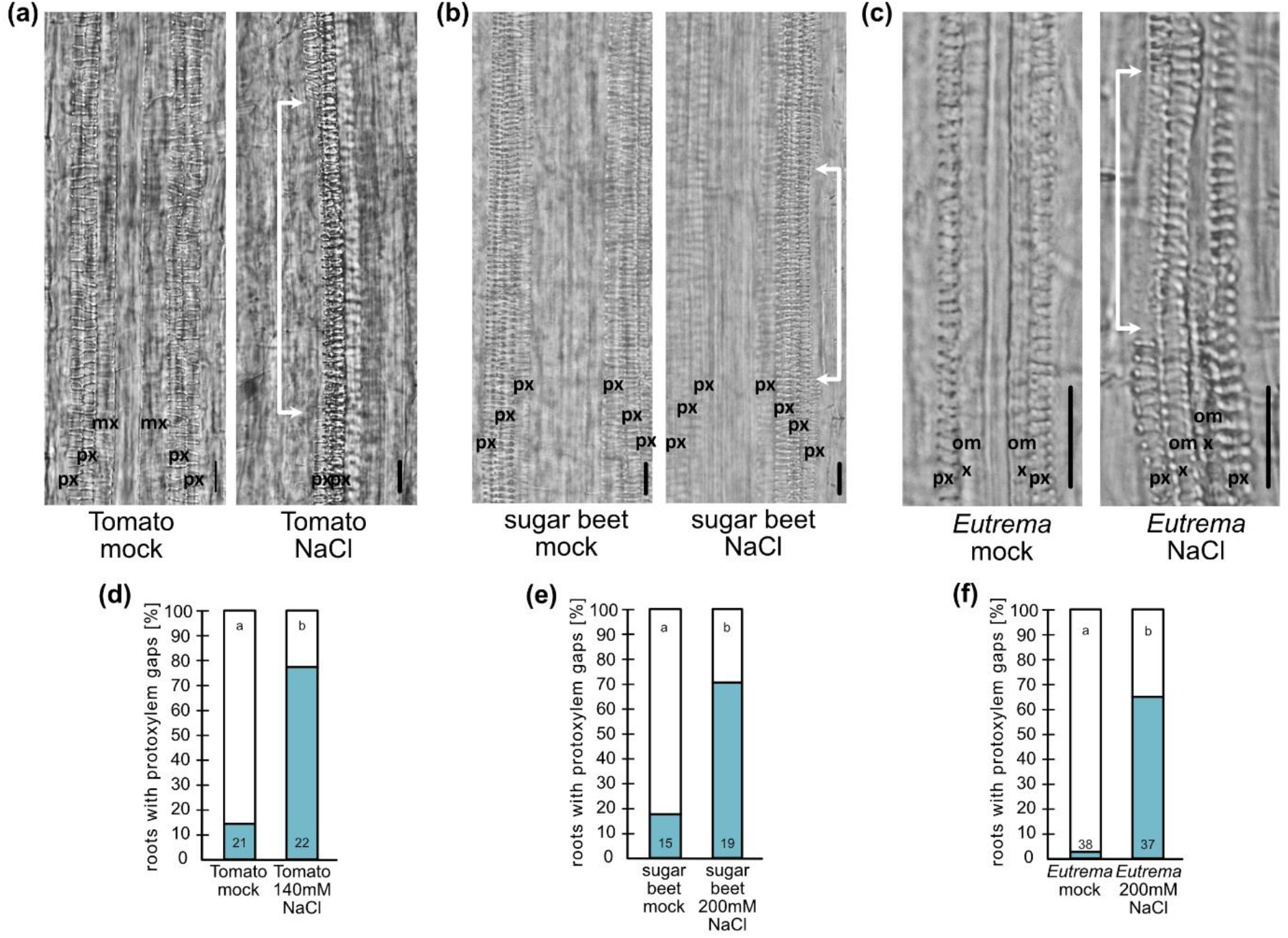
Protoxylem gaps are formed in several eudicot species upon salt. **(a)** Differential interference (DIC) images of tomato seedling root xylem after growth on NaCl or under mock conditions for 3 days. **(b)** Sugar beet seedling root xylem after growth on NaCl or under mock conditions for 3 days. **(c)** *Eutrema* seedling root xylem after growth on NaCl or under mock conditions for 3 days. White arrows indicate protoxylem gaps. px, protoxylem; mx, metaxylem. Scale bars, 50 μm. **(d)** Quantification of tomato roots exhibiting protoxylem gaps. **(e)** Quantification of sugar beet roots exhibiting protoxylem gaps. **(f)** Quantification of *Eutema* roots exhibiting protoxylem gaps. Numbers in the bars indicate n; letters indicate statistical significance with multiple Fisher’s exact test and BH correction, p < 0.05.

### Gibberellin levels affect protoxylem gap formation

Several studies have indicated that levels of bioactive GAs are reduced under saline conditions (Magome et al., 2008; Colebrook et al., 2014). We therefore tested if altered GA levels could affect protoxylem gap formation upon growth on salt in Arabidopsis. For this, we first grew plants on either 10 μM GA3, 1 μM GA4+7 or on paclobutrazol (PAC), which inhibits GA biosynthesis (Lee et al., 1985), with and without 140 mM NaCl. Growth on GA alone did not affect xylem development, but when we combined GA and salt we repeatedly noted a tendency of a lower number of plants forming protoxylem gaps, compared to salt-grown plants without GA added (Fig. S3a, b, c). PAC on its own, and in particular PAC together with salt, significantly enhanced the number of plants forming xylem gaps (Fig. 3a, Fig. S3d). These effects were not observed when PAC were added together with GA3, confirming that the effect of PAC on protoxylem gap formation both under control condition and on salt is due to its effect on GA levels (Fig. 3a, Fig. S3d). Consistent with these findings, the *ga4* mutant (Koornneef and van der Veen, 1980; Talon et al., 1990), defective in a late step in the synthesis of bioactive GA, displayed protoxylem gaps that could be restored by addition of GA3 (Fig. 3b, Fig. S3e). Upon growth on salt, a significantly higher frequency of the mutant plants formed protoxylem gaps compared to wildtype (Fig. 3b, Fig. S3e). Similarly, *ga1-3* and *ga1-5*, defective in an earlier GA-biosynthesis step (Sun et al., 1992), displayed increased frequency of protoxylem gap formation upon growth on salt, although not when grown under control conditions (Fig. S3f, g). Hence, these findings suggest that reduction in GA levels is linked with the protoxylem gap phenotype in Arabidopsis.

**Fig. 3:**
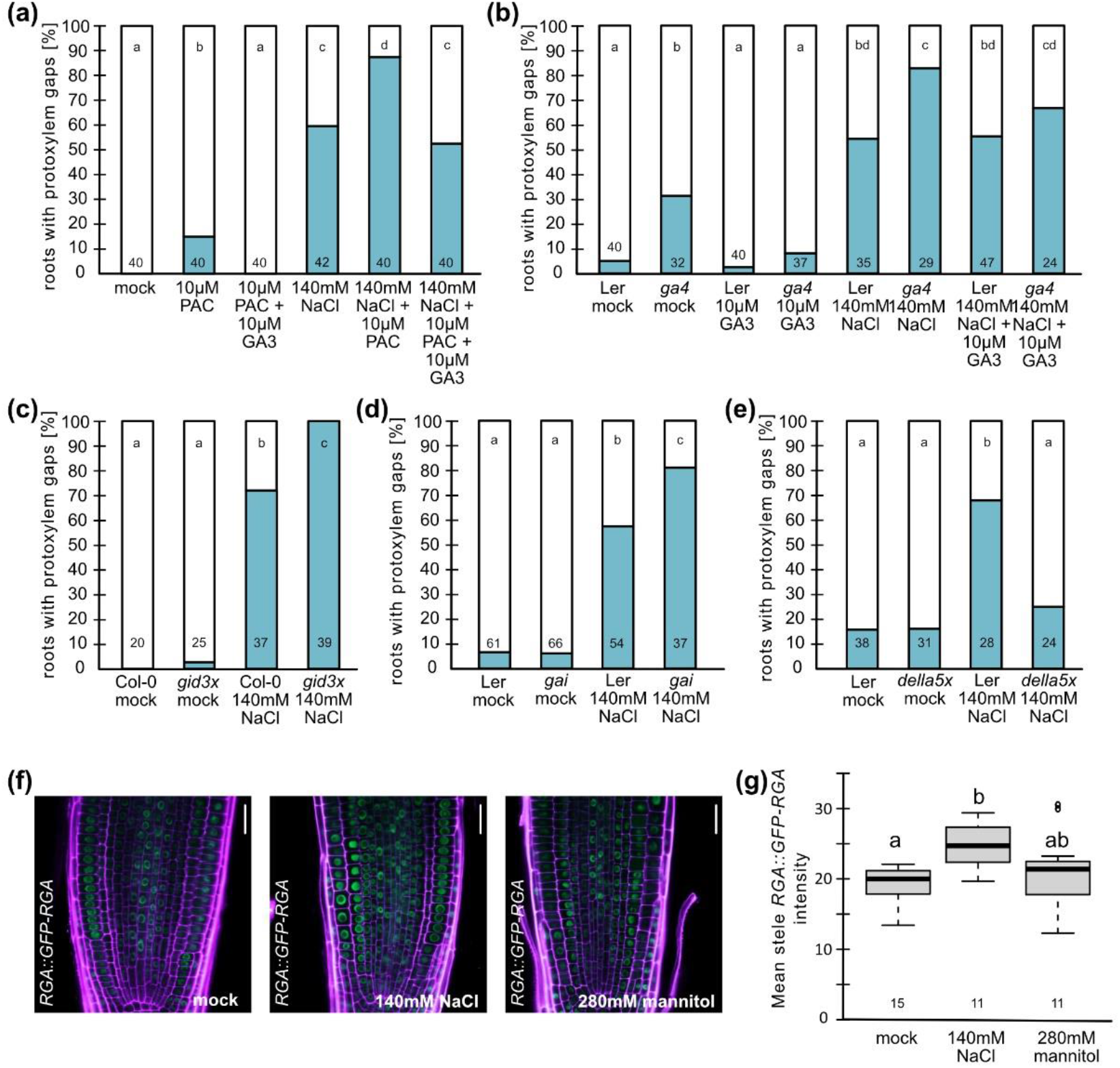
Reduced GA-levels and signaling induce protoxylem gap formation. **(a - e)** Quantifications of roots exhibiting protoxylem gaps of 6-day old Arabidopsis seedlings of the indicated genotypes grown under indicated conditions for 3 days. *Della5x* is *gai-t6 rga-t2 rgl1-1 rgl2-1 rgl3-4*. Numbers in bars are n; letters show statistical significance with multiple Fisher’s exact test and BH correction, p < 0.05. **(f)** Confocal micrographs of *RGA::GFP-RGA* in root meristems of 5-day old Arabidopsis seedlings after 6-9 h on 140mM NaCl, 280mM mannitol or mock conditions. Purple, PI-stain; Green, GFP. Scale bar, 20μm. **(g)** Quantification of mean stele *RGA::GFP-RGA* intensity. Numbers indicate n, letters statistical significance with Two-way ANOVA, p < 0.05.

### DELLAs are required for protoxylem gap formation upon salt

GA is sensed by GID-receptors, and consistent with our findings on altered GA levels the *gid1a-2 gid1b-3 gid1c-1* triple mutant, defective in GA perception (Griffiths et al., 2006), displayed both increased frequency of plants with protoxylem gap formation and of gaps per root when grown on salt (Fig. 3c, Fig. S3h). Upon GA perception, DELLAs, which act as transcriptional co-regulators, become degraded to allow GA responses (Locascio et al., 2013). The DELLA-protein GAI is stabilized in the *gai* mutant and thus acts as a constitutive GA-signalling repressor (Koorneef et al., 1985). Consistent with the involvement of DELLA-mediated signalling for xylem gap formation this mutant displayed enhanced frequency of xylem gap forming plants upon salt (Fig. 3d). Previously, it was shown that the two DELLA proteins, RGL3 and RGA, are stabilized upon exposure to salt (Achard et al., 2006; Geng et al., 2013; Shi et al., 2017). To analyse if growth on salt may affect RGA accumulation also in the stele of the root meristem, we grew the *RGA::GFP-RGA* translational reporter lines on 140mM NaCl or 280mM mannitol and analysed GFP signal intensity in procambium, xylem precursor cells and pericycle. This revealed that salt, but not mannitol, significantly enhanced RGA accumulation in the stele (Fig. 3f, g), consistent with the previous study.

The five DELLAs appeared to function redundantly in the control of xylem gap formation. The single *della* mutants *gai-td1* (Sessions et al., 2002), *rgl3-5* and *rga-28* (Tyler et al., 2004) had no effect on gap formation upon salt stress (Fig. S3i, j), while higher order mutants such as *rgl3-5 rga-28* and the *gai-t6 rga-24 rgl1-1 rgl2-1* (*della4x*) mutant (Ragni et al., 2011), lead to a gradual increase in the suppression of protoxylem gap formation (Fig. S3k, l). The DELLA quintuple (*della5x) gai-t6 rga-t2 rgl1-1 rgl2-1 rgl3-4* mutant, defective in all five DELLA genes (Koini et al., 2009), did not form significantly more protoxylem gaps upon salt stress than under mock condition (Fig. 3e). This suggests that protoxylem gap formation induced by high salinity requires multiple DELLA proteins.

### DELLA-activated VND6 contributes to protoxylem gap formation upon salt stress

To further understand the processes regulated by altered GA levels and DELLA-proteins in roots under salt-stress, we analyzed global gene expression changes in roots of 5-day old wildtype, *della5x*, *gai* and *ga4* seedlings after exposure to 140mM NaCl for 1h and/or 8h compared to control conditions (Fig. 4a, S4d). The time points were selected to show the response at the initial stress response phase, and when the plants had acclimatized and resumed growth and development (Geng et al., 2013). Corroborating our results suggesting that salt reduced levels of active GAs in the roots we found several GA-2-oxidases, which inactivates bioactive GAs, upregulated upon exposure to salt in the initial stress response phase (Table S1). As salt affects protoxylem differentiation, we focused our attention to genes previously found to be expressed in immature xylem in two different single cell transcriptomes (Denyer et al., 2019; Wendrich et al., 2020). Consistent with the notion that a subset of the early stress responding are under DELLA regulation we found gene ontologies (GOs) such as “response to stress”, “response to stimulus” and “response to other cellular components” enriched among genes that were upregulated in wild type and differentially expressed in the *della5x* mutant, among the xylem active genes (Table S2). Genes downregulated in wildtype upon salt exposure, differentially regulated by *della5x* and expressing in immature xylem were enriched for processes such as “genes encoding enzymes involved in hemicellulose synthesis” (Fig. S4f, g). Hence, cell wall modifications may happen rapidly, but we expected most of the factors underlying gap formation to primarily express at the acclimation phase at 8h. At the 8h time point 2887 genes were activated after salt exposure in wildtype (log2 fold change (FC) >0.5/ log2 FC <-0.5, padj < 0.05). Of these 450 had a reported xylem expression, of which 184 were differentially expressed in *della5x*, *ga4* or *gai* (p < 0.05) (Fig. 4b). These genes were enriched in GOs related to ‘response to stress’ but also ‘cell wall organization and biosynthesis’ (Fig. 4c), while corresponding down-regulated genes included e.g. ‘stress response’, ‘water transport’, ‘cellular processes’. As the cell walls are altered in the protoxylem gap cells, we focused our attention on the ‘cell wall organization and biosynthesis’ group of genes (Fig. 4d). Among these were several encoding proteins with functions relating to cellulose and hemicellulose biosynthesis along with the xylem master regulator VND6 as well as several alpha expansins. The xylem expressed list with differential expression in GA-mutants included also the previously described VND6 target *MYB DOMAIN PROTEIN 83* (*MYB83*) (McCarthy et al., 2009; Fig. 4d). While these genes were activated by salt, the *della5x* mutant displayed less strong activation and/or expression was further upregulated in *gai* or *ga4*. This is in agreement with our finding that *della5x* suppressed xylem gap formation, and also indicates that one or more of the DELLAs directly or indirectly activate transcription of these xylem regulator genes in wildtype under salt stress.

**Fig. 4:**
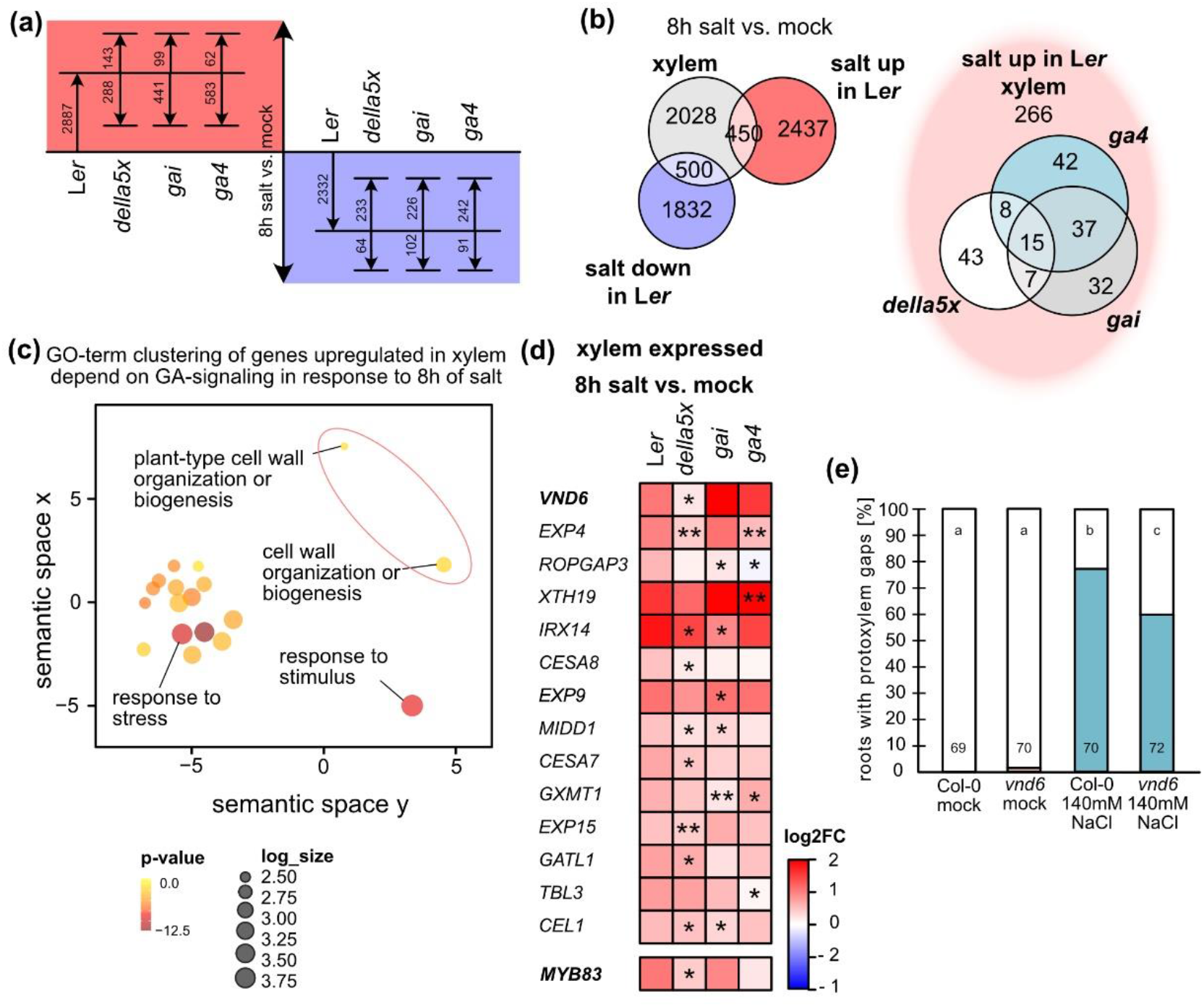
Cell wall related genes are differentially expressed in a DELLA-dependent manner upon salt. **(a)** Differentially expressed genes (log2 FC < −0.5/ >0.5, padj < 0.05) after 8h of salt exposure in L*er* (wildtype) roots and if indicated genotypes are significantly higher or lower expressed than L*er* (p < 0.05). *Della5x* is *gai-t6 rga-t2 rgl1-1 rgl2-1 rgl3-4*. **(b)** Left, Venn diagram of genes differentially expressed, up or down, in L*er* after 8h of salt exposure and genes expressed in xylem according to published single cell data sets (Denyer et al., 2019; Wendrich et al., 2020). Right, Venn diagram of xylem expressed genes upregulated by salt and differentially expressed (up or down) in *della5x*, *ga4* and/or *gai*. **(c)** REVIGO clustering (Supek et al., 2011) of GO-terms enriched among genes upregulated by 8h salt exposure, xylem expressed and differentially expressed in *della5x*, *gai* and/or *ga4*. **(d).** Heatmap of genes with differential expression relative L*er* within the GO-term clusters related to cell wall organization or biogenesis from (E), as well as *MYB83* which is in the dataset, but not highlighted in the GOs. Bold indicate master regulators of xylem development; *, p < 0.05; **, padj < 0.05. **(e)** Quantification of Col-0 (wildtype) and *vnd6* roots exhibiting protoxylem gaps. Numbers indicate n; letters statistical significance with multiple Fisher’s exact test and BH correction, p < 0.05.

To examine the potential influence of VND6 on xylem development upon salt stress, we analyzed the *vnd6* mutant (Kubo et al., 2005). Although it did not exhibit any apparent differences in xylem development under control conditions, significantly less protoxylem gaps were formed in *vnd6* upon salt stress compared to wild type (Fig. 4e), suggesting a role for VND6 in salt induced protoxylem gap formation. However, the gap reduction in *vnd6* was relatively limited, indicating that additional factors contribute to the xylem gap formation under salt.

### Expansins may be involved in DELLA dependent xylem gap formation

Xylem gaps are not only observed upon salt stress, but certain genotypes display gaps also under normal conditions, including protoxylem gaps in *ga4* (Fig. 3b) and metaxylem gaps upon inhibition of ABA signaling in the endodermis (Ramachandran et al., 2021). In search for a common set of genes differentially expressed under these conditions, we identified three genes: *EXP1*, *XYLOGLUCAN ENDOTRANSGLUCOSYLASE/ HYDROLASE* (*XTH20*) and a peroxidase family gene (*At2G18150*) (Fig. 5a, b). Neither *EXP1* nor *XTH20* are found in the immature xylem single cell transcriptomes (Denyer et al., 2019; Wendrich et al., 2020), but expression data from (Geng et al., 2013) indicates that both are upregulated in the stele in response to salt (Fig. S5a). As several genes encoding expansins were upregulated in response to salt in our analyses (Fig. 4d, S5b) and expansins previously have been related to salt stress tolerance and vascular development (Jadamba et al., 2020) we focused on EXP1. While the *expa1-1* mutant did not display any xylem deviations under normal conditions, this mutant could suppress protoxylem gaps formation upon salt stress (Fig. 5c), linking also EXP1 to modifications in xylem development upon salt stress.

**Fig. 5:**
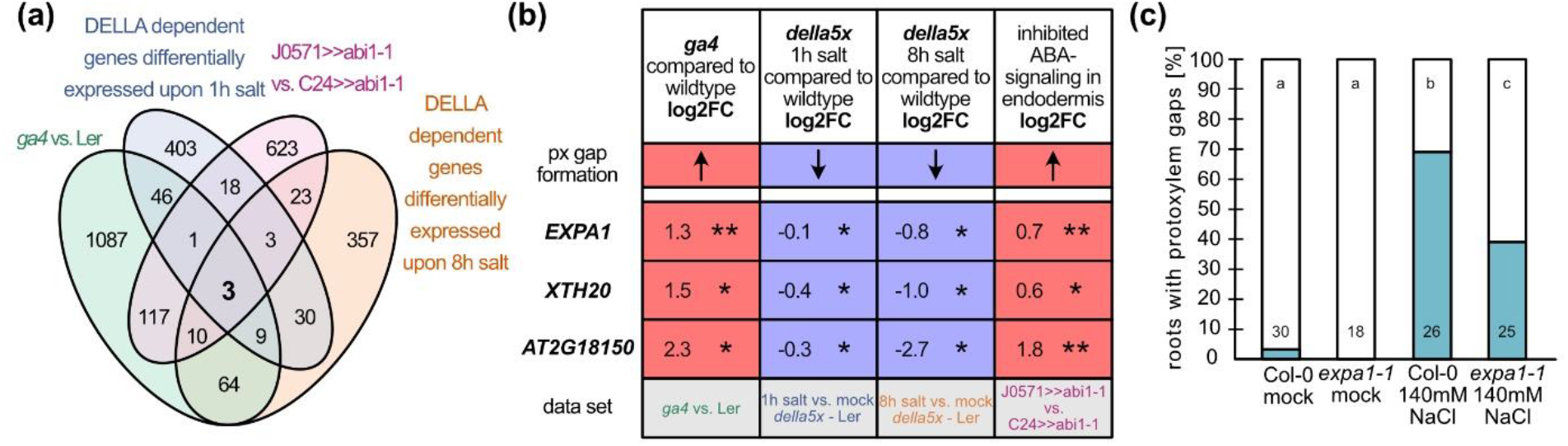
Cell wall modifying enzymes in xylem gap formation. **(a)** Venn diagram showing the overlap between different xylem gap related datasets: *ga4* vs. L*er* (this study, log2FC <-0.5/>0.5, p < 0.05), DELLA dependent genes differentially expressed upon 1h salt (this study; L*er* log2FC >0.5/-<0.5, padj < 0.05; *della5x* (*gai-t6 rga-t2 rgl1-1 rgl2-1 rgl3-4*) p < 0.05), DELLA dependent genes differentially expressed upon 8h salt (this study; L*er* log2FC >0.5/-<0.5, padj < 0.05; *della5x* p < 0.05), and *J0571*>>*abi1-1* vs. *C24*>>*abi1-1* (Ramachandran et al., 2021; log2FC >0.5/-<0.5, p < 0.05). **(b)** Differential expression relative to wildtype (log2FC) of genes shared by the four datasets, shown in (a). *, p < 0.05; **, padj < 0.05. **(c)** Quantification of roots exhibiting protoxylem gaps of 6-day old Arabidopsis seedlings of Col-0 (wildtype) and *expa1-1* upon growth on 140mM NaCl or under mock conditions for 3 days. Numbers indicate n; letters indicate statistical significance with multiple Fisher’s exact test and BH correction, p < 0.05.

Taken together, these results connect the xylem developmental regulator VND6, along with several of its well-known targets including genes encoding SCW-modifying enzymes, together with one or more alpha-expansins as GA-regulated factors important for the formation of protoxylem gaps under salt stress.

### Xylem gap formation correlates with better survival under salt stress

Next, we asked if the xylem gaps might help the plant to withstand salt stress. To this end, we compared the performance of mutants that produce less protoxylem gaps upon salt stress with mutants that produce more gaps. For mutants producing fewer gaps we chose *della5x* and *expa1-1* (Fig. 3e, 5c), and for mutants with excessive gap formation we selected *ahp6-1*, which affects protoxylem specification by interfering with the auxin-cytokinin balance (Mähönen et al., 2006) and a multiple *vnd*-mutant, *vnd1 vnd 2 vnd3 vnd7* (*vnd1237*), affecting xylem differentiation (Tan et al., 2018; Ramachandran et al., 2021; Fig. S6a, b). We grew 3-day old seedlings of these lines along with wild type on media containing 200 mM NaCl for 4 days, and then scored survival by the colour of the cotyledons (white versus green or pale green). The survival rate of the *della5x* mutant was significantly worse than wild type upon salt stress, and the *expa1-1* mutant displayed a similar trend (Fig. 6a, c, S6d, f). In contrast, both *ahp6-1* and *vnd1237* survived significantly better upon salt stress compared to wild type (Fig. 6a, b, S6d, e). When repeating the assay for *ahp6-1* with 7-day old plants, after onset of secondary growth (Smetana et al., 2019), we did not find a difference in salt stress tolerance compared to wild type, indicating that protoxylem gaps may be important mainly for young seedlings, prior to secondary growth initiation (Fig. S6c).

**Fig. 6:**
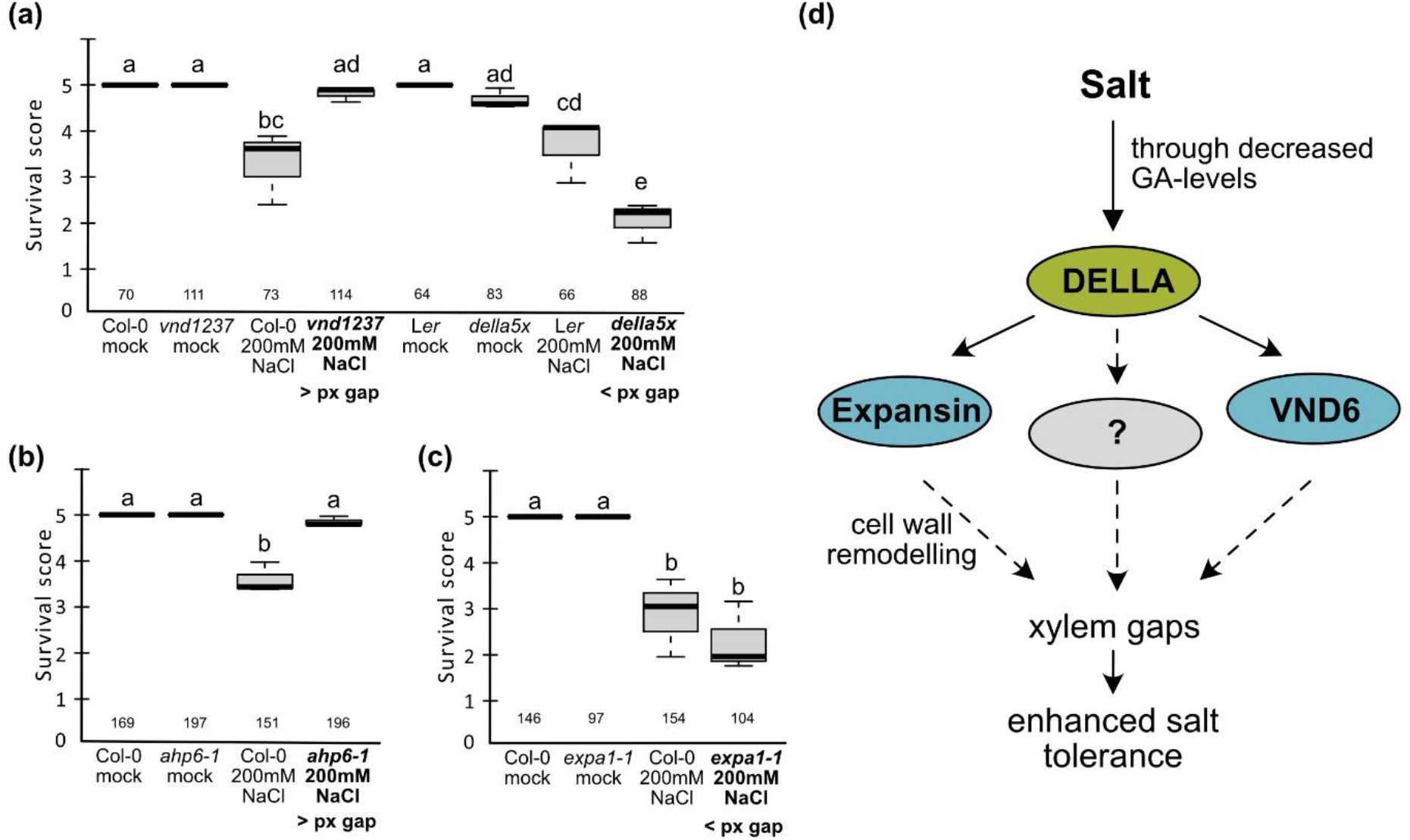
Enhanced protoxylem gap formation confers increased salt tolerance. **(a - c)** Survival after growth on 200mM NaCl or under mock conditions for 4 days for mutants with more (*vnd1 vnd2 vnd3 vnd7*, *vnd1237* and *ahp6-1*) or less (*della5x*, *gai-t6 rga-t2 rgl1-1 rgl2-1 rgl3-4*, and *expa1-1*) protoxylem (px) gap formation in response to salt stress, compared to wildtype (Col-0 or L*er*). Survival score was calculated by assigning plants with white cotyledons a score of 1, pale green 3 and green 5. These scores were multiplied and then divided by n analyzed plants. Numbers indicate n from three replicates. Letters indicate statistical significance with Two-way ANOVA, p < 0.05. **(d)** Proposed model for the molecular regulation of xylem gap formation in response to salt stress. Salt causes a decrease in levels of bioactive GAs (Achard et al., 2006) leading to the stabilization of DELLA proteins. One or more of the DELLAs promote expression of the xylem development master regulator *VND6*, along with genes for SCW formation, expansins and other cell wall remodeling genes, resulting in the occasional prevention of fully differentiated protoxylem cells, visible as protoxylem gaps, by an as of yet unknown mechanism. These gaps promote enhanced salt tolerance in young seedlings.

## Discussion

### Protoxylem gaps may promote seedling survival under high salinity

High soil salinity strongly impairs crop productivity and is a major problem in agriculture (Shannon and Grieve, 1999). Although salt affects plants in all developmental stages, the seedling establishment phase may be particularly vulnerable, and genetic traits affecting performance of early life stages contribute strongly to selection and local adaptation (Postma and Ågren, 2016). Here, we provide results showing that the extent to which a young seedling produce protoxylem gaps correlates with salt tolerance. We cannot exclude that other parameters also influence salt tolerance, but supporting the relevance of protoxylem gaps, such gaps are induced by moderate salinity not only in wildtype Arabidopsis seedlings (Col-0 and L*er* ecotypes), but also in seedlings of other eudicot species. Previous studies have found that processes that inhibit the transport of salt ions to the shoot via the xylem promote salt tolerance (Møller et al., 2009; Jiang et al., 2012). This may be seen in relatively salt tolerant Arabidopsis accessions which respond to salt by developing smaller vessels and less well developed xylem compared to phloem in the stem, potentially reducing hydraulic conductivity (Sellami et al., 2019a). As contrast, the *acl5* mutant which develop extensive xylem is salt hypersensitive (Shinohara et al., 2019). The protoxylem gaps formed in response to salt stress may be particularly important during the seedling establishment phase, as older *ahp6* seedlings with excessive protoxylem gaps behaved similar to wildtype on salt. In the very young seedling, water transport would rely relatively more on the protoxylem while most transport would occur via metaxylem and secondary xylem vessels in later developmental stages, likely decreasing the relative relevance of the protoxylem strands.

### Protoxylem gap formation upon high salinity requires reduced GA-signaling

Previously, we and others have found that plants exposed to water limiting conditions or elevated ABA levels respond by forming additional protoxylem strands and metaxylem closer to the root tip (Ramachandran et al., 2018; Bloch et al., 2019; Ramachandran et al., 2021). Here we show that these ABA mediated responses also occur upon exposure to moderate salt stress, but only relatively late. A more rapid response is the formation of protoxylem gaps, which happens independently of ABA signaling. Instead, multiple lines of evidence point towards reduced GA levels and/or signaling as critical for the formation of protoxylem gaps in response to salt stress. In particular, a mutant defective in all five DELLA genes, and thus with de-repressed GA-signaling, did not form protoxylem gaps upon salt stress. Our results are in line with previous data which suggest that high levels of GA render plants more sensitive to salt, and that GA-2-oxidases, which normally are upregulated by salt, reduce GA-levels and protect against salt stress (Achard et al., 2006; Magome et al., 2008; Colebrook et al., 2014). Consequently, *della*-mutants are less salt tolerant, while stabilisation of RGL3, along with the auxin-signalling repressor IAA17, confers salt stress resistance (Achard et al., 2006; Achard et al., 2008; Shi et al., 2017). While multiple studies connect GA-signaling with various steps of xylem development (Ashraf et al., 2002; Colebrook et al., 2014; Guo et al., 2015; Yamazaki et al., 2018; Singh et al., 2019), it has not been clear that GA’s effects on xylem morphology and differentiation impacts salt tolerance. Hence, our findings here explain at least partly how altered GA levels and signaling influence salt tolerance in the young seedling.

### Xylem master regulator VND6 is employed in gap formation under salt stress

A relatively large set of previously identified xylem-active genes were upregulated in a DELLA dependent manner under salt, including genes encoding the xylem differentiation master regulators VND6 and MYB83 (Kubo et al., 2005; McCarthy et al., 2009), along with multiple genes known to act downstream of these regulators encoding e.g. SCW cellulose synthases. This was a puzzling observation, as the protoxylem gaps had reduced SCW differentiation. However, the *vnd6* mutant displayed reduced capacity for gap formation suggesting that the activation of *VND6* upon salt indeed is connected to the intermittently inhibited protoxylem cell differentiation. Thus, these results suggest that *VND6* activated under salt may have a different and additional role than governing mextaxylem differentiation, and instead contribute to the modification of protoxylem differentiation. How VND6 orchestrate this feat is currently unknown. Intriguingly, several genes that were activated under salt stress, but less so in the *della5x* mutant, including *VND6*, displayed instead elevated expression in *della5x* compared to wild type under control conditions (Fig S4c, Table S1). This suggests that DELLAs repress *VND6* under control conditions, but activate it upon salt stress. It is known that DELLAs can act both as transcriptional activators and repressors depending on interaction partner (Locascio et al., 2013; Yoshida et al., 2014). Thus, identifying DELLA interacting partners under normal and high salt conditions will be important in future studies to elucidate how VND6 may shift activity to contribute to the promotion of protoxylem gaps under salt stress.

It is likely that other factors act in parallel with VND6, downstream of the DELLAs, as the *vnd6* mutant could not completely suppress gap formation under salt. We found that a mutant of *EXP1* also partially could repress xylem gap formation, indicating that cell wall remodeling also contributes to xylem gap formation. Previous studies have similarly found elevated expansin levels under salt stress, and cell walls may undergo extensive remodeling under salt acclimation (Shen et al., 2014). Altered SCW composition with a higher cellulose and hemicellulose content and reduced lignin was observed in the Arabidopsis stem after salt stress, resulting in enhanced vessel elasticity preventing collapse due to the osmotic stress (Sellami et al., 2019b). In rice, overexpression of *OsEXPA7* led to changes in the vasculature and increased salt tolerance (Jadamba et al., 2020). From our results, we cannot tell if cell wall modifications happen exclusively in the xylem or also in other tissues, and it will be relevant to further assess cell-specific effects upon salt stress.

### Conclusion

Salt stress induces local inhibition of protoxylem differentiation causing protoxylem gaps in young Arabidopsis seedlings. Gap formation requires DELLA-mediated repression of GA-signaling, and it is likely that salt stress triggers a reduction in bioactive GA-levels, thus stabilizing DELLA proteins. Under salt stress, DELLAs promote a set of VND6-regulated factors, including SCW-differentiation enzymes as well as other enzymes such as alpha-expansins as depicted in the model in Fig. 6d. Mutational analysis of VND6 and EXP1 link these factors specifically to the formation of protoxylem gaps under salt. These gaps may confer salt tolerance to young seedlings and thus provide an adaptive advantage. Seedlings of other distantly related eudicot species similarly from protoxylem gaps upon salt stress, suggesting evolutionary conservation of this trait.

## Acknowledgements

We thank C. Melnyk, Swedish University of Agricultural Sciences (SLU); L. Ragni, University of Tübingen; C. Köhler, Max Planck Institute, Potsdam, C. Dixelius, SLU and Maribo Hilleshög, and the Nottingham Arabidopsis Stock Centre for materials; M. Englund for technical assistance; A. Zhang for bioinformatics advice and C. Melnyk and E. Sundberg for carefully reading the manuscript. We acknowledge support from the Nilsson Ehle Foundation (F.A.) and Formas (2017-00857, A.C.). Part of the computations were enabled by resources in projects SNIC 2021/22-647 and SNIC 2021/23-547 provided by the Swedish National Infrastructure for Computing (SNIC) through Uppsala Multidisciplinary Centre for Advanced Computational Science (UPPMAX) partially supported by the Swedish Research Council through grant agreement no. 2018-05973.

## Author contributions

Design of the research, F.A., A.C.; Performance of the research, F.A.; Data analysis, F.A, A.C; Writing – Original Draft, F.A.; Writing – Review & Editing, F.A., A.C.; Funding Acquisition, F.A., A.C.

## Data availability

All data and material produced in this study are available on request from the corresponding author.

## Declaration of interests

The authors declare no competing interests.

## Supplemental Information

Fig. S1: Protoxylem gaps are formed in response to salt

Fig. S2: Protoxylem gaps are formed in several eudicot species upon salt

Fig. S3: Reduced GA-levels and signaling induce protoxylem gap formation

Fig. S4: Cell wall related genes are differentially expressed in a DELLA-dependent manner upon salt

Fig. S5: Cell wall modifying enzymes in xylem gap formation

Fig. S6: Enhanced protoxylem gap formation confers increased salt tolerance

Table S1: Differentially expressed genes in roots of L*er*, *della5x*, *gai* and *ga4* upon growth on salt for 1h and 8h

Table S2: Enriched GOs from Panther analysis of xylem expressed genes up- and downregulated by 1h/ 8h of salt exposure in DELLA-/ GA-dependent manner

Table S3: Summary of statistical analyses

Methods S1: Key resources used in this study

## Notes

### Competing Interest Statement

The authors have declared no competing interest.

