## Supplementary material for "Salinity induces discontinuous protoxylem by a DELLA-dependent mechanism promoting salt tolerance in Arabidopsis seedlings": Fig_S1-6_Table_S2

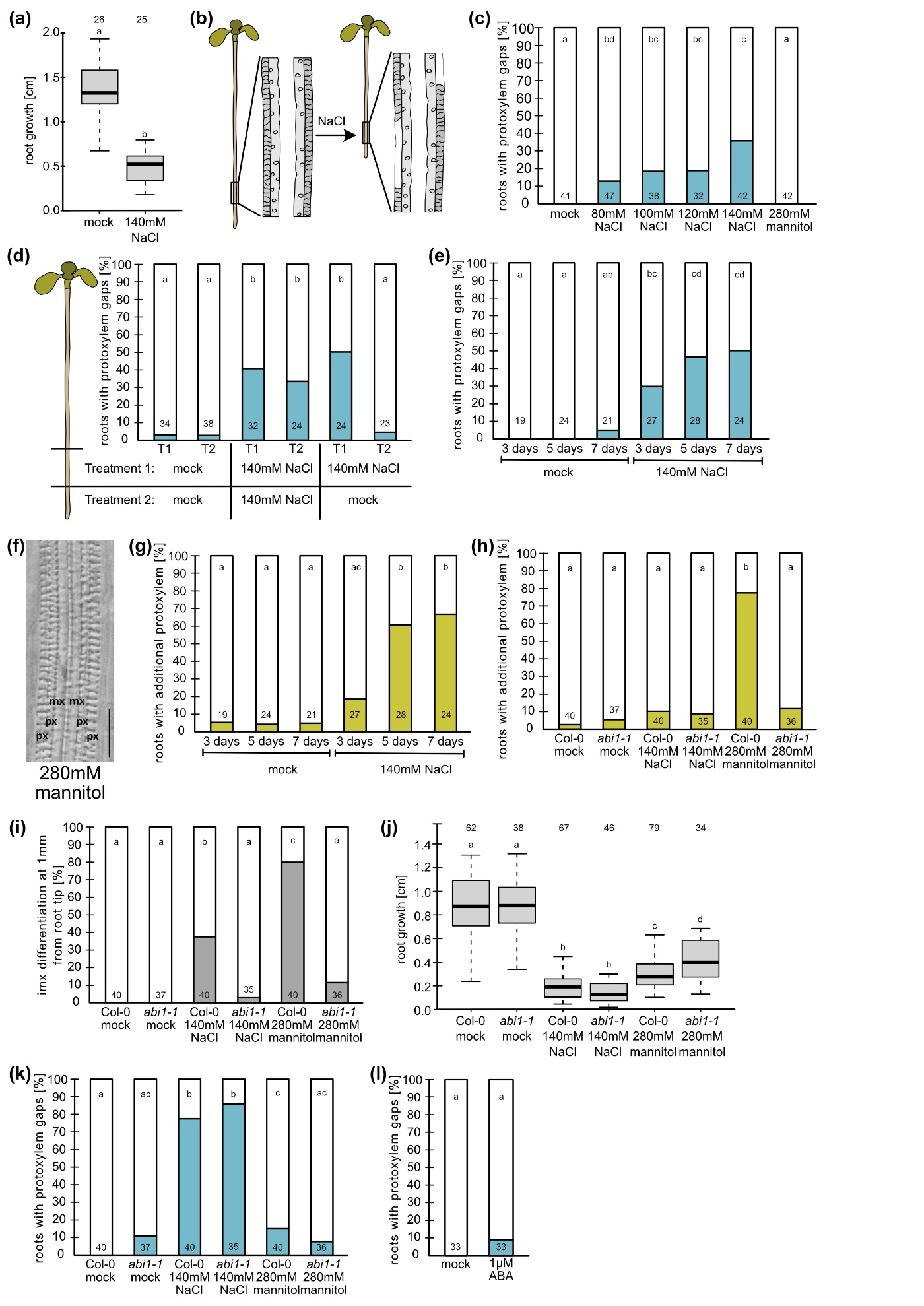

**Fig. S1: Protoxylem gaps are formed in response to salt.** All data are from roots of 6-day old Arabidopsis seedlings grown for 3 days on NaCl, mannitol, ABA or under mock conditions, unless otherwise stated. **(a)** Root growth of Arabidopsis seedlings on NaCl or mock plates. Numbers above the graph indicate n roots analyzed; letters indicate statistical significance with t-test, p < 0.05. **(b)** Cartoon of protoxylem gap formation along the root upon exposure to salt. **(c)** Quantification of number of roots with protoxylem gaps after growth on different concentrations of salt or mannitol, where 280 mM mannitol is iso-osmolaric to 140 mM NaCl. **(d)** Seedlings were transferred for three days to Treatment 1 (T1) and then transferred to Treatment 2 (T2) for three days. The parts of the roots that were grown under the different treatments were collected and analyzed separately. **(e)** Quantification of roots with protoxylem gaps after growth on salt for 3, 5 or 7 days. **(f** **-** **g)** Quantification of roots with extra protoxylem after growth on salt for 3, 5 or 7 days. **(h)** Quantification of extra protoxylem in wild type (Col-0) or *abi1-1* roots after growth on salt or mannitol. **(i)** Quantification of roots exhibiting differentiated inner metaxylem (imx) at 1 mm from the root tip, after growth on salt or mannitol. **(j)** Root growth of Col-0 or *abi1-1* on salt or mannitol. Letters indicate statistical difference with Two-way-ANOVA; p < 0.05. **(k)** Quantification of number of roots with protoxylem gaps in Col-0 and *abi1-1* after growth on salt or mannitol. **(l)** Quantification of number of roots with protoxylem gaps in Col-0 after growth on mock or 1µM ABA. In **c-e**, **g-i** and **k-l,** numbers in bars indicate n; letters indicate statistical significance with multiple Fisher’s exact test and BH correction, p < 0.05.

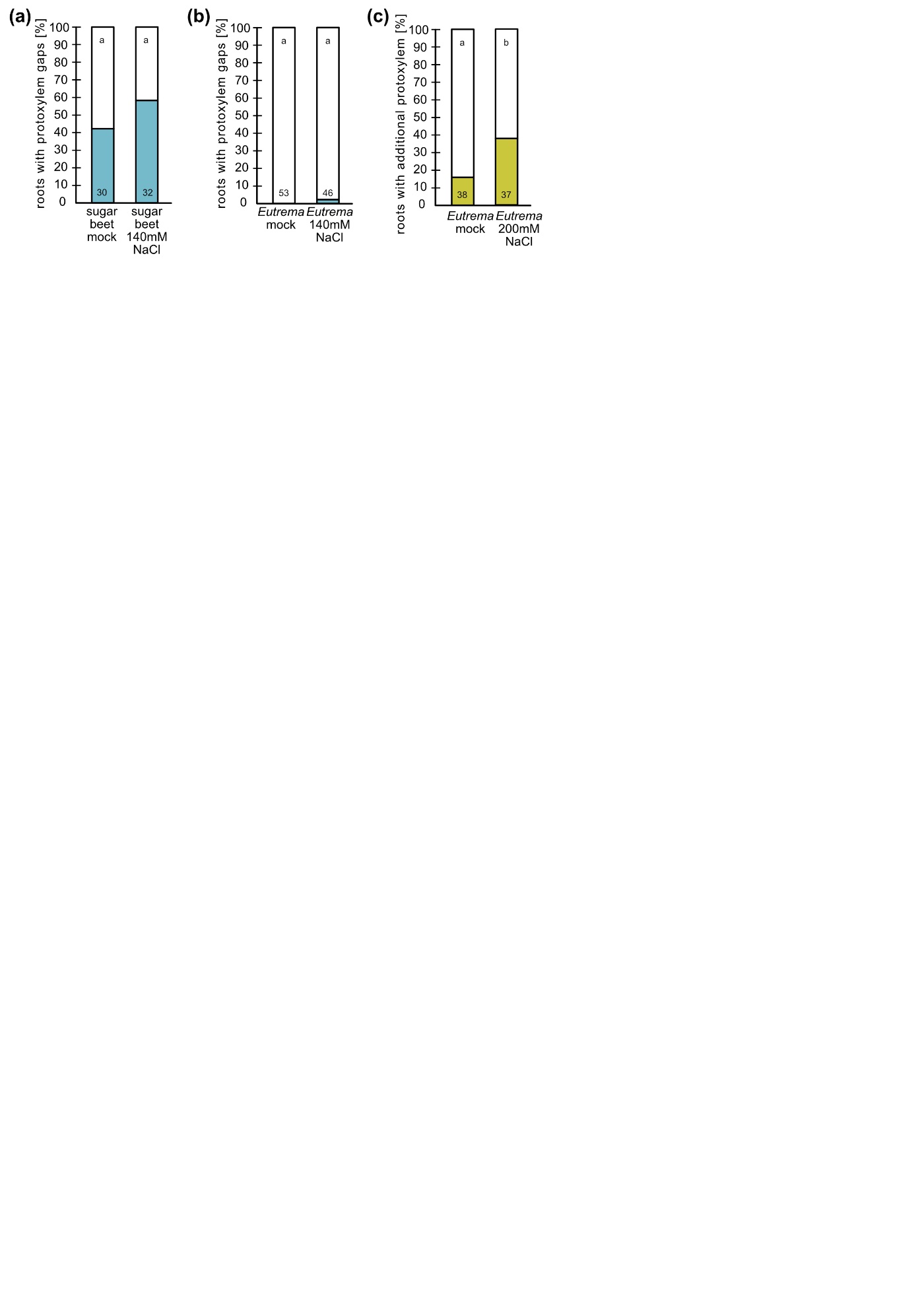

**Fig. S2: Protoxylem gaps are formed in several eudicot species upon salt. (a)** Quantification of sugar beet roots exhibiting protoxylem gaps after growth for 3 days on 140 mM NaCl. **(b)** Quantification of *Eutrema* roots exhibiting protoxylem gaps after growth for 3 days on 140 mM NaCl. **(c)** Quantification of *Eutrema* roots exhibiting extra protoxylem after growth for 3 days on 200 mM NaCl. Numbers in bars indicate n; letters indicate statistical significance with multiple Fisher’s exact test and BH correction, p < 0.05.

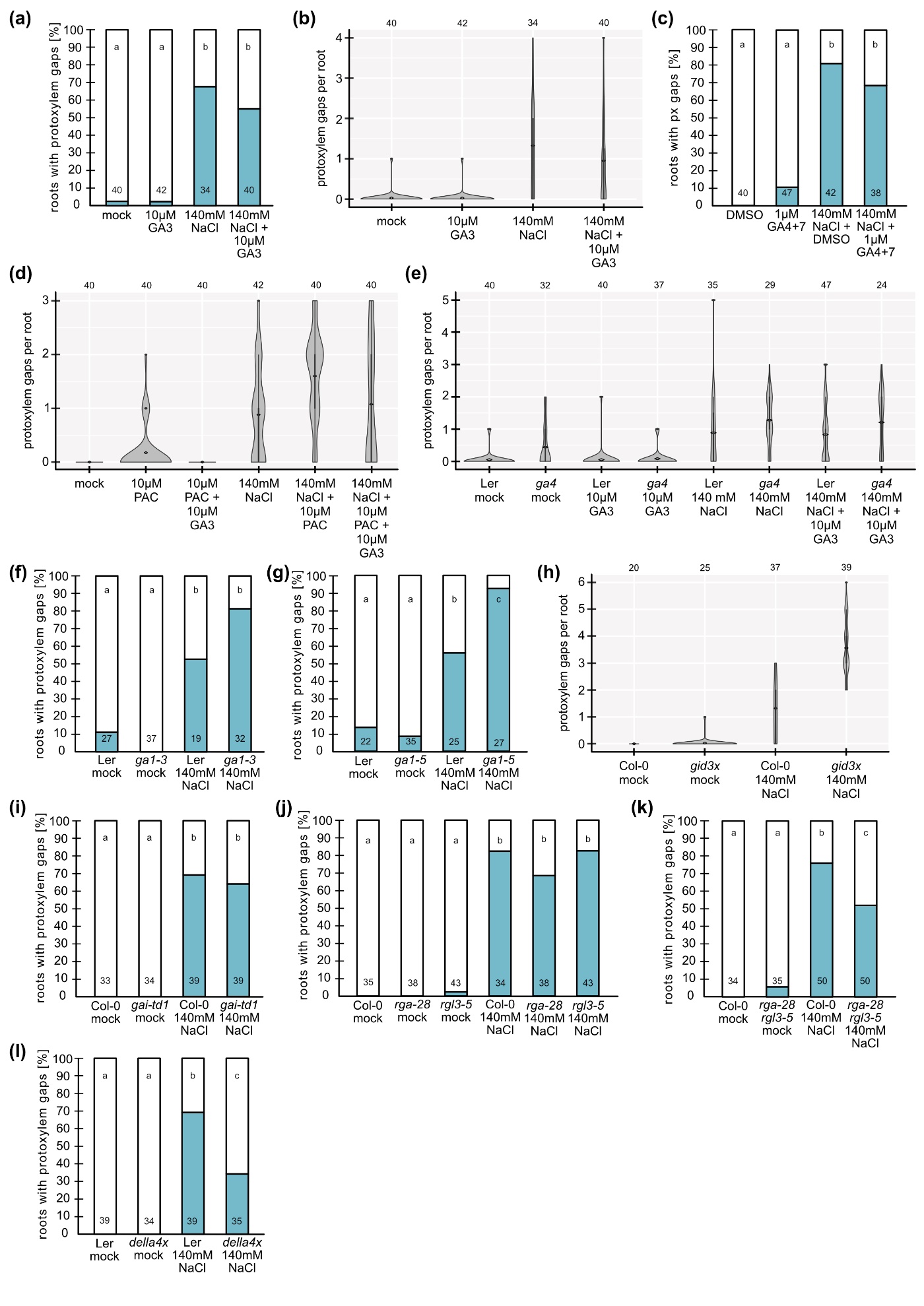

**Fig. S3: Reduced GA-levels and signaling induce protoxylem gap formation.** All results presented were observed in roots of 6-day old Arabidopsis seedlings of indicated genotypes grown for 3 days on 140mM NaCl or under mock conditions, and with indicated treatments of GA3, GA4+7 (dissolved in DMSO) or Paclobutrazol (PAC). *Della4x* is *gai-t6 rga-24 rgl1-1 rgl2-1*. **(a), (c), (f - g), (i - l)** Quantification of roots exhibiting protoxylem gaps. Numbers indicate n; letters indicate statistical significance with multiple Fisher’s exact test and BH correction, p < 0.05. **(b)**, **(d)**, **(e)**, **(h)** Violin plots displaying number of protoxylem gaps per root. Numbers above the graph indicate n.

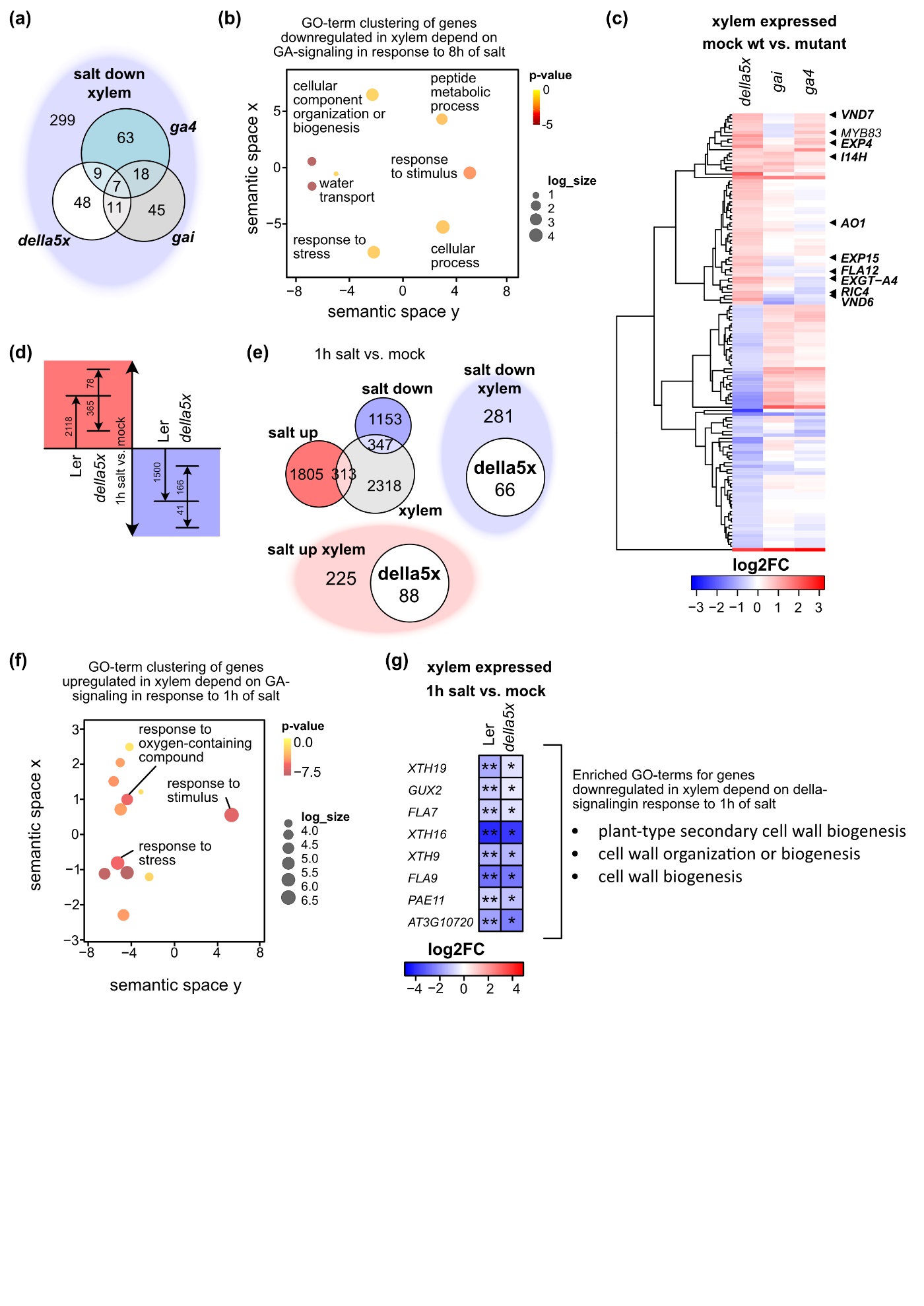

**Fig. S4: Cell wall related genes are differentially expressed in a DELLA-dependent manner upon salt. (a)** Venn diagram-section (light blue) of genes downregulated in wildtype and expressed in xylem according to published single cell data sets (Denyer et al., 2019; Wendrich et al., 2020), see fig. 4B. The fraction of genes differentially expressed in *della5x* (*gai-t6 rga-t2 rgl1-1 rgl2-1 rgl3-4*), *ga4* and/or *gai* is displayed on top of the section. **(b)** REVIGO clustering (Supek et al., 2011) of GO-terms enriched for genes that are downregulated upon salt, xylem expressed and differentially expressed in *della5x*, *gai* and/or *ga4*. **(c)** Heatmap of genes that are differentially regulated in della5x vs. wildtype under mock (log2FC < -0.5/ > 0.5, p < 0.05), and filtered for xylem-expression. **(d)** Differentially expressed genes (log2FC < -0.5/ >0.5, padj < 0.05) after 1h of salt exposure in Ler (wildtype) and the fraction of those genes differentially expressed in della5x (p < 0.05). **(e)** Venn diagram of genes that are up- and downregulated in wildtype after 1h of salt exposure and genes expressed in xylem according to published single cell data sets (Denyer et al., 2019; Wendrich et al., 2020). Venn diagram fractions display upregulated (pink) or downregulated (light blue) xylem expressed genes differentially expressed in della5x. **(f)** REVIGO clustering (Supek et al., 2011) of GO-terms enriched for genes that are upregulated upon 1h salt, xylem expressed and differentially expressed in della5x. **(g)** Heatmap of genes extracted from GO-term enrichment analysis for genes that are downregulated upon 1h salt. Only the three shown GO-terms were enriched. *, p < 0.05; **, padj < 0.05.

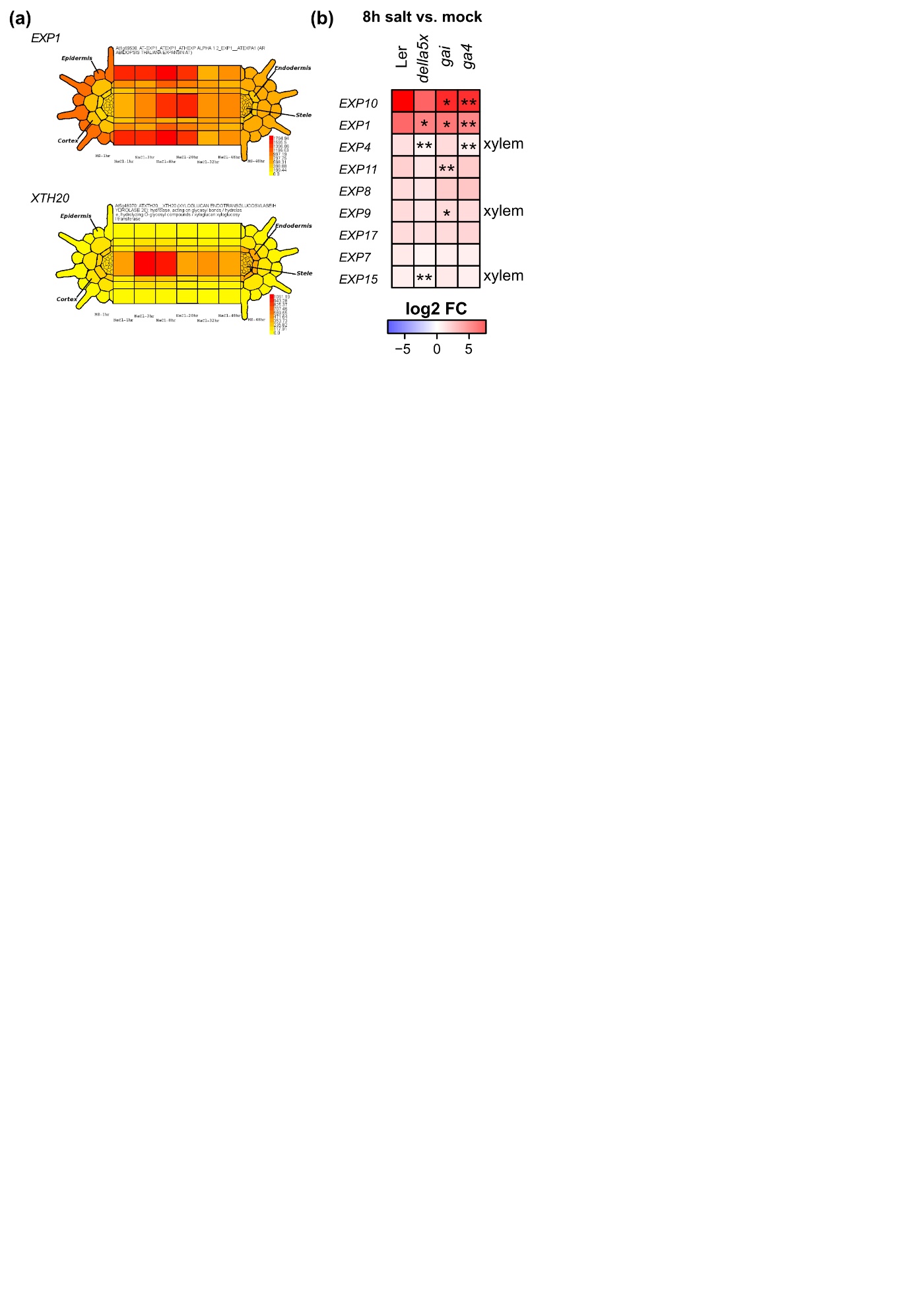

**Fig. S5: Cell wall modifying enzymes in xylem gap formation. (a)** Tissue specific expression of *EXP1* and *XTH20* during a time course of salt exposure extracted from the eFP browser (Geng et al., 2013). **(b)** Heatmap showing expression of alpha-expansins after 8h of salt exposure in L*er* (wild type), *della5x* (*gai-t6 rga-t2 rgl1-1 rgl2-1 rgl3-4*), *gai* and *ga4* *, p < 0.05; **, padj < 0.05 (see also Supplementary Table 1).

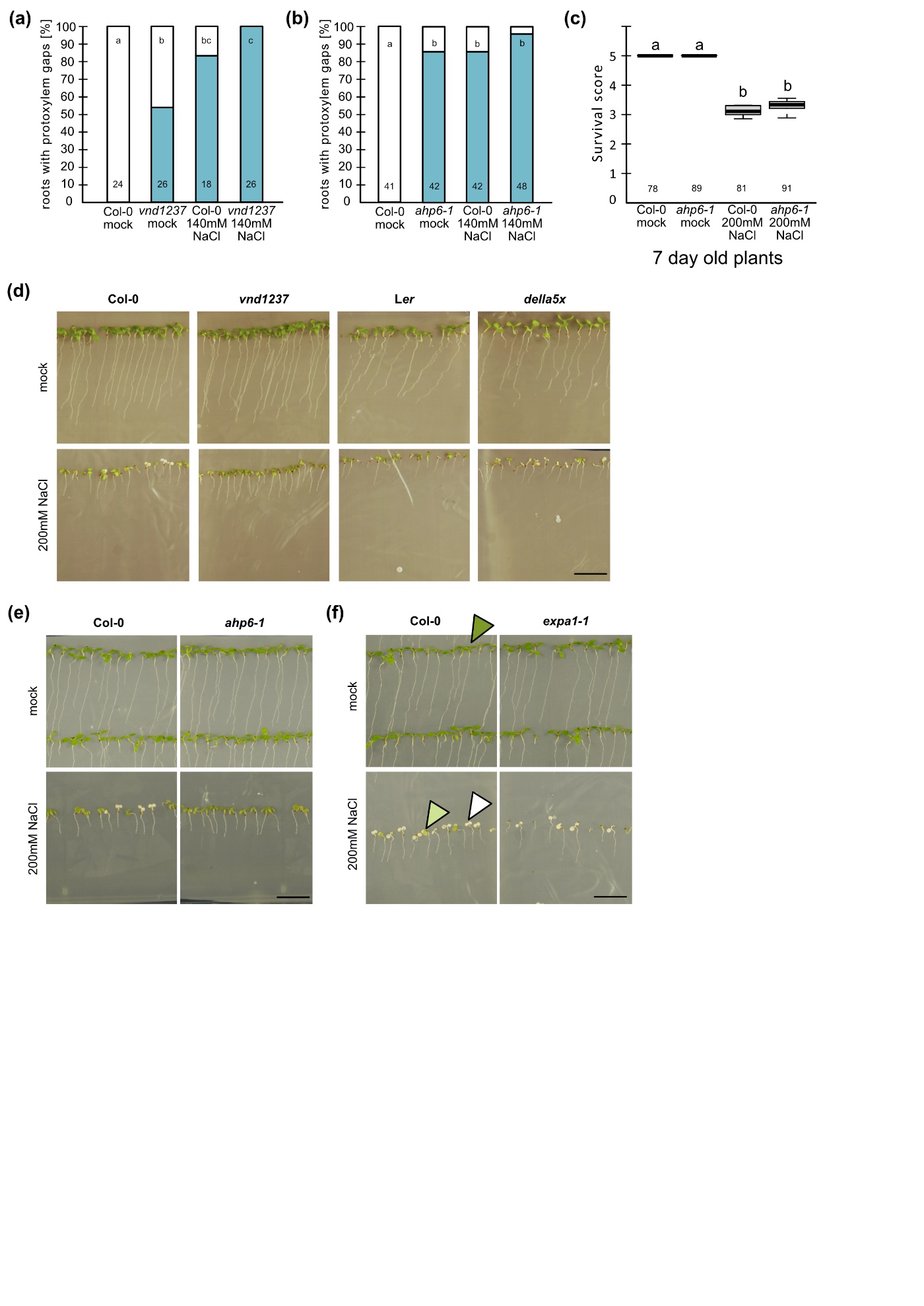

**Fig. S6: Enhanced protoxylem gap formation confers increased salt tolerance. (a)** Quantification of roots exhibiting protoxylem gaps in 6-day old Arabidopsis Col-0 or *vnd1 vnd2 vnd3 vnd7* (*vnd1237*) grown for 3 days on 140mM NaCl or under mock conditions. Numbers indicate n; letters indicate statistical significance with multiple Fisher’s exact test and BH correction, p < 0.05. (**b**) Quantification of roots exhibiting protoxylem gaps in 6-day old Arabidopsis Col-0 or *ahp6-1* grown for 3 days on 140mM NaCl or under mock conditions. **(c)** Salt tolerance assay after growth on 200mM NaCl or mock conditions for 4 days for 7 day old *ahp6-1* seedling. Survival score was calculated by assigning plants with white cotyledons a score of 1, pale green 3 and green 5. These scores were multiplied and then divided by n analyzed plants. Numbers indicate n from five replicates. Letters indicate statistical significance with Two-way ANOVA, p < 0.05. **(d, e)** Pictures of seedlings of indicated genotypes, grown for 4 days on mock or 200mM NaCl. *della5x* is *gai-t6 rga-t2 rgl1-1 rgl2-1 rgl3-4*. Scale bar, 1 cm. **(f)** Pictures of seedlings of indicated genotypes, grown for 4 days on mock or 200mM NaCl. Arrowheads point at seedlings with green, pale green and white cotyledons. Scale bar, 1 cm.

**Table S1:** **Differentially expressed genes in roots of L*er*, *della5x*, *gai* and *ga4* upon growth on salt for 1h and 8h, related to Fig. 4, 5, S4, S5**

**(a)** and **(b)** RNA-Seq on roots (1 cm) of L*er* and *della5x* (*gai-t6 rga-t2 rgl1-1 rgl2-1 rgl3-*4) treated for 1h with 140mM NaCl or mock. **(a)** Differential expression analysis evaluating the impact of the mutant background on the effect of 140mM NaCl treatment (combinatorial effect). Log2 fold-changes (log2FC) were extracted from the pairwise comparison of mock-treatment for each genotype, while p values and adjusted p (padj) were extracted from the pairwise comparison between mutants and wildtype. P values and padj for L*er* were extracted from the pairwise comparison with mock treatment. Genes upregulated by NaCl in L*er* with log2FC>0.5 are marked in green lettering, and those with log2FC<-0.5 in red, genes with padj<0.05 are marked in bold, genes with p<0.05 in italics. Genes significantly upregulated upon NaCl in L*er* (padj<0.05), which are significantly reduced in mutants (p<0.05) are marked in blue lettering. **(b)** Differential expression analysis between L*er* and *della5x*. Log2FC, p values and padj were extracted from pairwise comparisons. Genes with log2FC>0.5 in mutant compared to L*er* are marked in green lettering, and those with log2FC<-0.5 in red. Genes with padj<0.05 are marked in bold; genes with p<0.05 in italics. **(c)** and **(d)** RNA-Seq on roots (1 cm) of L*er*, *della5x, gai, ga4* treated for 8h with 140mM NaCl or mock. **(c)** Differential expression analysis evaluating the impact of the mutant background on the effect of 140mM NaCl treatment (combinatorial effect). Log2FC were extracted from the pairwise comparison of mock-treatment for each genotype, while p values and padj were extracted from the pairwise comparison between mutants and wildtype. P values and padj for L*er* were extracted from the pairwise comparison with mock treatment. Genes upregulated by NaCl in L*er* with log2FC>0.5 are marked in green lettering, and those with log2FC<-0.5 in red, genes with padj<0.05 are marked in bold, genes with p<0.05 in italics. Genes significantly upregulated upon NaCl in L*er* (padj<0.05), which are significantly reduced in mutants (p<0.05) are marked in blue lettering. **(d)** Differential expression analysis between L*er* and *della5x*, *gai*, and/or *ga4*. Log2FC, p values and padj were extracted from pairwise comparisons. Genes with log2FC>0.5 in mutant compared to L*er* are marked in green lettering, and those with log2FC<-0.5 in red. Genes with padj<0.05 are marked in bold; genes with p<0.05 in italics. Xylem_Denyer2019: genes expressed in the immature xylem identified by scRNASeq (Denyer et al., 2019); Xylem_Wendrich2020: genes expressed in the immature xylem identified by scRNASeq (Wendrich et al., 2020).

**Table S2:** Enriched GOs from Panther analysis of xylem expressed genes up- and downregulated by 1h/ 8h of salt exposure in DELLA-/ GA-dependent manner

| **Xylem expressed genes upregulated by 1h of salt exposure in a DELLA-dependent manner** | | |
| --- | --- | --- |
| **GO biological process complete** | | **P value** |
| GO:0009737 | response to abscisic acid | 0.00000836 |
| GO:0033993 | response to lipid | 0.00000382 |
| GO:0010033 | response to organic substance | 0.00000935 |
| GO:0042221 | response to chemical | 4.47E-10 |
| GO:0050896 | response to stimulus | 3.08E-08 |
| GO:0009725 | response to hormone | 0.00000468 |
| GO:0009719 | response to endogenous stimulus | 0.00000678 |
| GO:0097305 | response to alcohol | 0.00000983 |
| GO:1901700 | response to oxygen-containing compound | 0.000000119 |
| GO:0009414 | response to water deprivation | 0.0147 |
| GO:0006950 | response to stress | 0.000000135 |
| GO:0009415 | response to water | 0.0182 |
| GO:0001101 | response to acid chemical | 0.0323 |
| GO:0009628 | response to abiotic stimulus | 1.18E-09 |
| GO:0006970 | response to osmotic stress | 0.00355 |
| GO:0009266 | response to temperature stimulus | 0.0121 |
| **Xylem expressed genes downregulated by 1h of salt exposure in a DELLA-dependent manner** | | |
| **GO biological process complete** | | **P value** |
| GO:0009834 | plant-type secondary cell wall biogenesis | 0.019 |
| GO:0071554 | cell wall organization or biogenesis | 0.0108 |
| GO:0042546 | cell wall biogenesis | 0.0153 |
| **Xylem expressed genes upregulated by 8h of salt exposure in a GA-dependent manner** | | |
| **GO biological process complete** | | **P value** |
| GO:0009414 | response to water deprivation | 0.000000201 |
| GO:0006950 | response to stress | 1.34E-10 |
| GO:0050896 | response to stimulus | 5.31E-10 |
| GO:0009415 | response to water | 0.00000031 |
| GO:1901700 | response to oxygen-containing compound | 0.000000389 |
| GO:0042221 | response to chemical | 2.65E-13 |
| GO:0010035 | response to inorganic substance | 0.0000297 |
| GO:0001101 | response to acid chemical | 0.00000102 |
| GO:0009628 | response to abiotic stimulus | 0.0000137 |
| GO:0009636 | response to toxic substance | 0.0284 |
| GO:0009737 | response to abscisic acid | 0.000000889 |
| GO:0033993 | response to lipid | 0.0000164 |
| GO:0010033 | response to organic substance | 0.000144 |
| GO:0009725 | response to hormone | 0.0000284 |
| GO:0009719 | response to endogenous stimulus | 0.0000456 |
| GO:0097305 | response to alcohol | 0.00000112 |
| GO:0071669 | plant-type cell wall organization or biogenesis | 0.0126 |
| GO:0071554 | cell wall organization or biogenesis | 0.00552 |
| GO:0006970 | response to osmotic stress | 0.00167 |
| GO:0009605 | response to external stimulus | 0.000128 |
| **Xylem expressed genes downregulated by 8h of salt exposure in a GA-dependent manner** | | |
| **GO biological process complete** | | **P value** |
| GO:0080170 | hydrogen peroxide transmembrane transport | 0.0413 |
| GO:0009987 | cellular process | 0.0139 |
| GO:0006833 | water transport | 0.00000408 |
| GO:0042044 | fluid transport | 0.00000408 |
| GO:0006412 | translation | 0.0297 |
| GO:0043043 | peptide biosynthetic process | 0.0328 |
| GO:0006518 | peptide metabolic process | 0.0109 |
| GO:0071840 | cellular component organization or biogenesis | 0.035 |
| GO:0006950 | response to stress | 0.0241 |
| GO:0050896 | response to stimulus | 0.000956 |

**Table S3:** Summary of statistical analyses

**Methods S1:** Key resources used in this study

| **REAGENT or RESOURCE** | **SOURCE** | **IDENTIFIER** |
| --- | --- | --- |
| **Chemicals, peptides, and recombinant proteins** | | |
| Murashige and Skoog Medium (MS) | Duchefa Biochemie | Cat#M0222.0050 |
| MES monohydrate | Duchefa Biochemie | Cat#M1503.0250 |
| Bactoagar | Swab | Cat#B1000-1 |
| Abscisic acid (ABA) | Sigma | Cat#14375-45-2 |
| Gibberellic acid 3 | Duchefa Biochemie | Cat#G0907 |
| Gibberellic acid 4+7 | Duchefa Biochemie | Cat#G0938 |
| Paclobutrazol (PAC) | Sigma | Cat#46046 |
| Mannitol | VWR Chemicals | Cat#25311.366 |
| Chloralhydrate | Sigma | Cat#15307 |
| Urea | Sigma | Cat#57-13-6 |
| Sodium deoxycholate | Sigma | Cat#1065040250 |
| Xylitol | Sigma | Cat#X3375 |
| Propidium iodide | Sigma | Cat#P4170 |
| **Critical commercial assays** | | |
| RNeasy Plant Mini Kit | Qiagen | Cat#74904 |
| Qubit BR RNA Assay | Invitrogen | Cat#Q10211 |
| **Deposited data** | | |
| Raw and processed RNAseq data files | This study | GEO: GSE195912 |
| Raw and processed RNAseq data files | (Ramachandran et al., 2021) | GEO: GSE169367 |
| **Experimental models: organisms/strains** | | |
| *Arabidopsis thaliana*: Col-0 | Widely distributed | N/A |
| *Arabidopsis thaliana:* L*er* | Widely distributed | N/A |
| *Solanum lycopersicum* cv. Moneymaker | Nelson garden and Plantagen | N/A |
| *Beta vulgaris* cv. Davinci | Maribo Hilleshög/ Christina Dixelius, SLU | N/A |
| *Eutrema salsugineum* | Claudia Köhler, Max Planck Inst. Potsdam | N/A |
| *Arabidopsis thaliana*: *abi1-1C* | (Kanno et al., 2012) | N/A |
| *Arabidopsis thaliana*: *vnd1 vnd2 vnd3 vnd7* in Col-0 background | (Ramachandran et al., 2021) | N/A |
| *Arabidopsis thaliana:* *gai-t6 rga-t2 rgl1-1 rgl2-1 rgl3-4* in L*er* background | (Koini et al., 2009) | N/A |
| *Arabidopsis thaliana: gai-t6 rga-24 rgl1-1 rgl2-1* in L*er* background | (Ragni et al., 2011) | N/A |
| *Arabidopsis thaliana: rga-28* in Col-0 background | (Tyler et al., 2004) | N/A |
| *Arabidopsis thaliana: rgl-3* in Col-0 background | NASC seed stock center | N/A |
| *Arabidopsis thaliana: rga-28 rgl3-5* in Col-0 background | This study | N/A |
| *Arabidopsis thaliana: gai-td1* in Col-0 background | (Plackett et al., 2014) | N/A |
| *Arabidopsis thaliana: gai* in L*er* background | (Koorneef et al., 1985) | N/A |
| *Arabidopsis thaliana: gid1a-2 gid1b-3 gid1c-1* in Col-0 background | (Griffiths et al., 2006) | N/A |
| *Arabidopsis thaliana: ga1-3* in L*er* background | (Sun et al., 1992) | N/A |
| *Arabidopsis thaliana: ga1-5* in L*er* background | (Sun et al., 1992) | N/A |
| *Arabidopsis thaliana: ga4* in L*er* background | (Koornneef and van der Veen, 1980; Talon et al., 1990) | N/A |
| *Arabidopsis thaliana: ahp6-1* in Col-0 background | (Mähönen et al., 2006) | N/A |
| *Arabidopsis thaliana: expa1-1* in Col-0 background | NASC seed stock center | N/A |
| *Arabidopsis thaliana*: *vnd6* in Col-0 background | (Kubo et al., 2005) | N/A |
| **Software and algorithms** | | |
| Zeiss Zen Black 2.3 SP1 | Zeiss | <https://www.zeiss.com/> |
| Zeiss Zen Blue 2.3 lite and 2.5 | Zeiss | <https://www.zeiss.com/> |
| R 4.02 and R studio 1.2.5019 | (R Core Team; RStudioTeam, 2019) | <https://www.r-project.org/>  <https://rstudio.com/> |
| Microsoft Excel 2016 | Microsoft | N/A |
| CellSet v1.5.1 | (Pound et al., 2012) | <https://sourceforge.net/projects/cellset/> |
| Affinity Designer 1.7 | Affinity | N/A |
| Fiji/Image J 2.0.0 Win64 or 2.0.0-rc-68/1.52h | (Schindelin et al., 2012) | <https://fiji.sc/> |
| Bioconductor 3.11 | Bioconductor (Huber et al., 2015) | <https://bioconductor.org/> |
| Fastp | (Chen et al., 2018) | <https://github.com/OpenGene/fastp> |
| MulitQC | (Ewels et al., 2016) | <https://multiqc.info/> |
| SortMeRNA | (Kopylova et al., 2012) | <https://bioinfo.lifl.fr/RNA/sortmerna/> |
| Trimmomatic | (Bolger et al., 2014) | <http://www.usadellab.org/cms/?page=trimmomatic> |
| FastQC | (Andrews, 2010) | <https://www.bioinformatics.babraham.ac.uk/projects/fastqc/> |
| HiSAT2 | (Kim et al., 2019) | <http://daehwankimlab.github.io/hisat2/> |
| HTSeq_Count | (Anders et al., 2015) | <https://htseq.readthedocs.io/en/release_0.9.1/count.html> |
| PANTHER 16.0 | (Mi et al., 2019; Mi et al., 2021) | <http://go.pantherdb.org/> |
| REVIGO | (Supek et al., 2011) | <http://revigo.irb.hr/> |
| **Other** | | |
| Zeiss LSM780 confocal microscope | Zeiss | <https://www.zeiss.com/> |
| Zeiss LSM800 confocal microscope | Zeiss | <https://www.zeiss.com/> |
| Zeiss Axioscope A1 | Zeiss | <https://www.zeiss.com/> |
